# Bloom syndrome helicase contributes to germ line development and longevity in zebrafish

**DOI:** 10.1101/2021.03.16.435627

**Authors:** Tamás Annus, Dalma Müller, Bálint Jezsó, György Ullaga, Gábor M. Harami, László Orbán, Mihály Kovács, Máté Varga

**Affiliations:** Department of Genetics, ELTE Eötvös Loránd University, Budapest, Hungary; Institute of Enzymology, Research Centre for Natural Sciences, Eötvös Loránd Research Network, Hungary; Department of Anatomy, Cell and Developmental Biology, ELTE Eötvös Loránd University, Budapest, Hungary; ELTE-MTA „Momentum” Motor Enzymology Research Group, Department of Biochemistry, ELTE Eötvös Loránd University, Budapest, Hungary; Frontline Fish Genomics Research Group, Department of Applied Fish Biology, Institute of Aquaculture and Environmental Safety, Hungarian University of Agriculture and Life Sciences, Georgikon Campus, Keszthely, Hungary; MTA-ELTE Motor Pharmacology Research Group, Department of Biochemistry, ELTE Eötvös Loránd University, Budapest, Hungary

**Keywords:** Bloom syndrome, BLM, RecQ, DNA repair, germ line, development, longevity, zebrafish

## Abstract

RecQ helicases - also known as the ‘guardians of the genome’ - play crucial roles in genome integrity maintenance through their involvement in various DNA metabolic pathways. Aside from being conserved from bacteria to vertebrates, their importance is also reflected in the fact that in humans impaired function of multiple RecQ helicase orthologs are known to cause severe sets of problems, including Bloom, Werner or Rothmund-Thomson syndromes. Our aim was to create and characterize a zebrafish (*Danio rerio*) disease model for Bloom syndrome, a recessive autosomal disorder. In humans, this syndrome is characterized by short stature, skin rashes, reduced fertility, increased risk of carcinogenesis and shortened life expectancy brought on by genomic instability. We show that zebrafish *blm* mutants recapitulate major hallmarks of the human disease, such as shortened lifespan and reduced fertility. Moreover, similarly to other factors involved in DNA repair, some functions of zebrafish Blm bear additional importance in germ line development, and consequently in sex differentiation. Unlike *fanc* genes and *rad51*, however, *blm* appears to effect its function independent of *tp53*. Therefore, our model will be a valuable tool for further understanding the developmental and molecular attributes of this rare disease, along with providing novel insights into the role of genome maintenance proteins in somatic DNA repair and fertility.

## Introduction

Originally described as a form of dwarfism accompanied by discolored skin of the nose and cheeks, Bloom’s Syndrome (BSyn) is a rare monogenic autosomal recessive disorder with less than 300 reported cases (Adam et al., 1993; Bloom, 1954). Symptoms of the disease include below average height and weight, lesions on exposed skin areas, reduced fertility and shortened life expectancy most often brought on by heightened proneness to cancer development (Cunniff et al., 2017). The defective gene responsible for the condition was mapped to 15q26.1 in the human genome and named Bloom (*BLM;* (German et al., 1994)). *BLM* encodes a 3’-5’ DNA helicase showing homology to the *E. coli* RecQ protein (Ellis et al., 1995; Karow et al., 1997). RecQ homologs are evolutionarily highly conserved proteins involved in a variety of genome integrity maintenance processes (Croteau et al., 2014).

While RecQ helicases are all involved in processes related to DNA metabolism and gene expression, human diseases linked to malfunctions of individual RecQ-family proteins lead to well-defined sets of symptoms, suggesting that their molecular functions are distinct. BLM in itself is required for precise double-stranded DNA break (DSB) repair, crossover patterning regulation, telomere maintenance, processing of DNA replication intermediates, and rDNA metabolism (Hatkevich et al., 2017; Schawalder et al., 2003; Seol et al., 2019).

Zebrafish is an attractive candidate to model RecQ family-related diseases as, despite an additional whole genome duplication event called teleost-specific genome duplication (TGD) in the common ancestor of teleosts (Christoffels et al., 2004; Taylor et al., 2003), all five human RecQ paralogs (*RECQL1, WRN, BLM, RECQL4* and *RECQL5*) have a single zebrafish ortholog each (Supplementary Figure 1A). In order to determine whether the species is indeed a viable model for BSyn, we generated a null allele for *blm* by CRISPR/Cas9 and analyzed its potential effects on lifespan, gonad differentation, fertility, histology, and DNA repair efficiency. As multiple genes involved in DSB can influence sex determination (SD) and differentiation (SDiff) in zebrafish (see below), we also tested the sex ratios of our mutants.

The specific mechanisms of zebrafish SD and SDiff in their full complexity are yet to be deciphered (for reviews see: (Kossack and Draper, 2019; Liew and Orbán, 2014; Orbán et al., 2009)). Unlike the ZZ/ZW chromosome-based process of wild and recently domesticated zebrafish strains (Wilson et al., 2014), SD of domesticated lines relies on a polygenic system with environmental factors playing secondary roles. This is presumably due to the loss of the original sex chromosomes during genome manipulations utilized in the domestication process. This indicates that SD not only differs from species to species in teleosts (Devlin and Nagahama, 2002; Penman and Piferrer, 2008; Wang et al., 2018), but it seems to vary among strains/lines in zebrafish (Anderson et al., 2012; Kossack and Draper, 2019; Liew and Orbán, 2014; Wilson et al., 2014). Over the last two decades, researchers have identified key components of sex differentiation of the species, shedding light on some of the processes involved. Interestingly, zebrafish genes primarily involved in genome maintenance have also been linked to sex differentiation. Knockout of genes associated with the Fanconi anemia (FA) pathway leads to complete or partial male bias, while masculinisation can be observed in homozygous mutants for *rad51* (Botthof et al., 2017; Ramanagoudr-Bhojappa et al., 2018; Rodríguez-Marí et al., 2010; Rodríguez-Marí et al., 2011; Shive et al., 2010).

Irrespective of their eventual sex, zebrafish larvae initially develop a ‘juvenile ovary’ giving rise to early oocytes (Takahashi1977, 1977; Wang et al., 2007). The direction of gonad development is contingent on the survival or apoptotic removal of these early oocytes. In developing males they undergo apoptosis, as the juvenile ovary transforms into testis (Uchida et al., 2002). Consistent with an anti-apoptotic role, elevated NF-κB signaling was shown to enhance the differentiation of females (Pradhan et al., 2012); furthermore, canonical Wnt signaling, a pathway involved in the SD of mammals as well, was shown to have similar effects (Pradhan et al., 2012; Sreenivasan et al., 2014; Vainio et al., 1999). An additional feature of zebrafish sex is its reliance on primordial germ cells (PGCs). These diploid stem cells are specified in the embryo and migrate into the gonadal ridge during the early development (for review see: (Marlow, 2015)). During gonad development, PGCs differentiate into gonadal stem cells (GSCs): ovarian stem cells (OSCs; (Hanna and Hennebold, 2014)) in females and spermatogonial stem cells (SSCs; (Kubota and Brinster, 2018)) in males. This is a general feature of gonad development of vertebrates and many multicellular invertebrates as well (Richardson and Lehmann, 2010).

The initially unimodal distribution of PGC numbers among individuals shifts to a bimodal one during early development in fishes due to their differential amplification between the two sexes: larvae with high and low PGC counts being prospective females and males, respectively (Saito et al., 2007). In zebrafish, experimental reduction of PGC count during early development leads to male bias, while its increase promotes female bias (Tzung et al., 2015a; Tzung et al., 2015b; Ye et al., 2019). As one would expect, PGC-ablated fish develop into sterile males, containing empty ‘testicular shells’ in place of their testes (Siegfried and Nüsslein-Volhard, 2008; Slanchev et al., 2005).

Although zebrafish do not undergo natural sex reversal as adults, the sexual plasticity of the gonads is maintained as shown by the fact that inhibition of estrogen synthesis in fertile adult females induces their masculinisation (Takatsu et al., 2013). This phenomenon taken together with the success of restoring male fertility by the transplantation of ovarian germ cells into sterile host larvae is consistent with the proposal that yet undifferentiated stem cells remain in the gonads of adult fish (Wong et al., 2011). In the absence of oocytes, female-to-male sex change occurs, thus oocytes are required for the maintenance of female sexual phenotype, most likely through their interaction with somatic cells expressing aromatase (Dranow et al., 2016; Dranow et al., 2013; Rodríguez-Marí and Postlethwait, 2011). As mentioned before, zebrafish homozygous for mutations of *fa* genes and *rad51* show partial or complete bias towards male development. This phenomenon could be attributed to increased germ cell apoptosis, as the all-male phenotypes could be rescued by the concurrent knockout of *tp53*, a central gene in apoptotic regulation (Aubrey et al., 2018; Botthof et al., 2017; Ramanagoudr-Bhojappa et al., 2018; Rodríguez-Marí et al., 2010).

Here we show that in zebrafish *blm* loss-of-function affects both somatic and germline cells, and Blm is necessary both during the early mitotic expansion of PGCs and during meiosis in males. Unexpectedly, the *blm* phenotype does not seem to be rescued by the impairment of *tp53*.

## Results

### Generation of blm loss-of-function zebrafish

The zebrafish genome contains a single *blm* ortholog on chromosome 18 (ENSDARG00000077089, GRCz11; Figure 1A). Similarly to many *fanc* genes and *rad51*, elevated levels of *blm* transcripts can be detected in the oocyte (Figure 1B,C). These maternal transcripts (many of them unpolyadenylated) undergo rapid degradation during mid-blastula transition and the zygotic expression of the gene starts at later stages of gastrulation (Figure 1B-D) (White et al., 2017; Winata et al., 2018).

**Figure 1:**
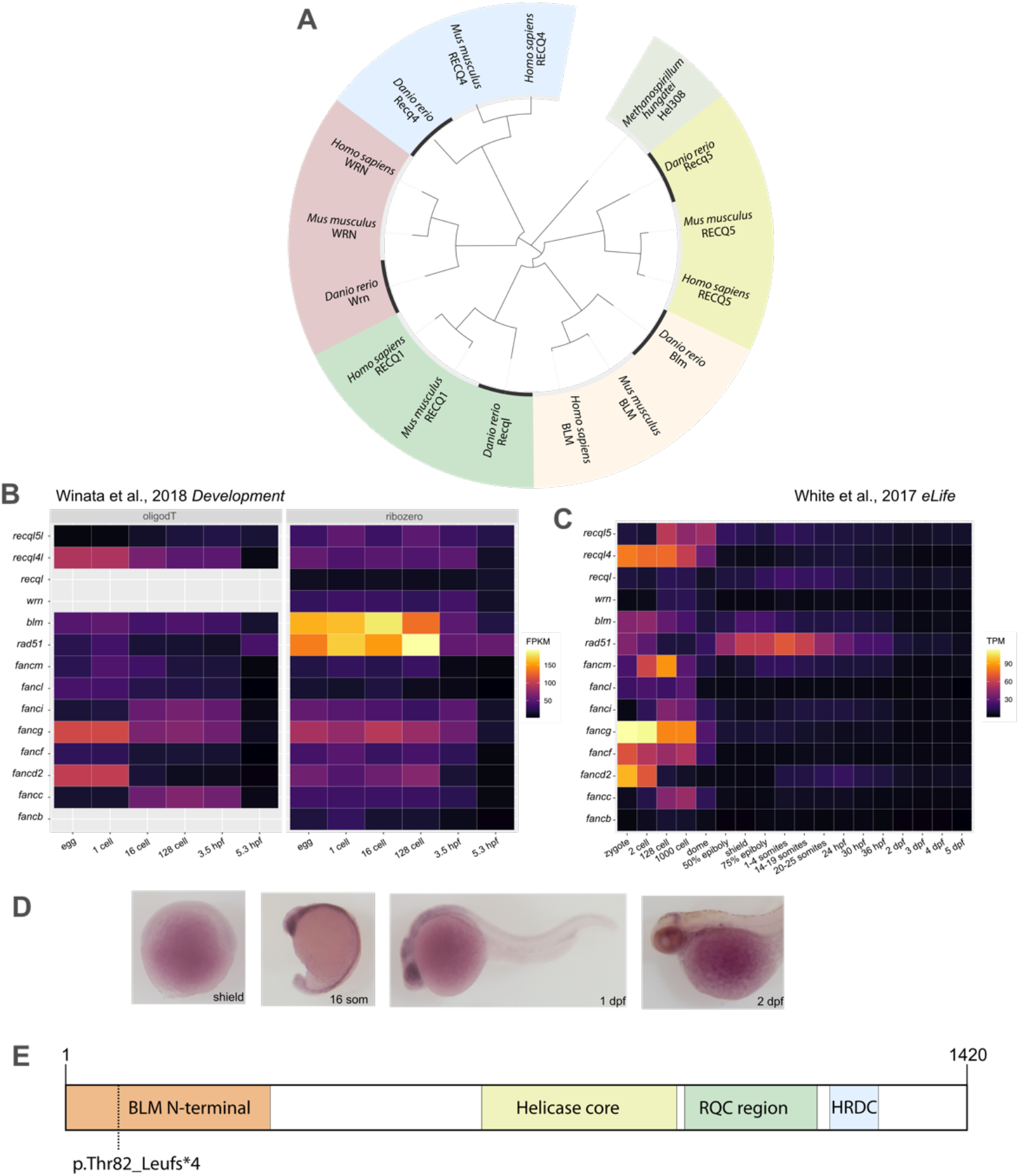
RecQ homologs in the zebrafish genome and their expression. (A) Phylogenetic relationships of the five RecQ homologs that can be identified in the zebrafish genome based on similarities between the helicase ATPase and helicase C-terminal domains. (*M. hungatei* Hel308 DNA helicase was used as an external reference for eukaryotic RecQs.) (B,C) The expression of genes related to double strand DNA breaks during zebrafish development. Fragments per kilobase of exon model per million reads mapped (FPKM) and transcripts per kilobase million (TPM) values of the different genes at given stages as shown by the datasets in the respective papers (White et al., 2017; Winata et al., 2018). (D) Spatial distribution of *blm* RNA during early stages of development detected by whole mount *in situ* hybridization. (E) Schematic domain structure of the zebrafish Blm and the position of the p.Thr101Leufs*4 mutation.

In order to gain a better understanding of the role of Blm in zebrafish we created a mutant allele using CRISPR/Cas9 by targeting the fourth exon of the gene. We were able to isolate a small indel (c.301_304delACAAinsTT) that results in a premature stop codon (p.Thr101Leufs*4) and a severely truncated protein (Figure 1E).

### Blm loss-of-function does not impair early somatic development and DNA damage sensitivity of zebrafish

The overall development and viability of *blm*^−/−^ zebrafish embryos and larvae was not compromised under normal conditions, as they were present in mendelian ratios in *blm*^+/−^ incross progenies (see below). Immunostaining for the phosphorylated H2A.X histone variant γ-H2AX, a marker of double stranded DNA breaks (DSBs) did not reveal differences between 2 days post-fertilisation (dpf) *blm*^−/−^ embryos and their siblings (Figure 2A,B). This result is strikingly different to the stark increase in γ-H2AX punctae observed in *rad51* mutants (Botthof et al., 2017), despite the involvement of both proteins in homologous recombination (HR)-based DNA break repair.

**Figure 2:**
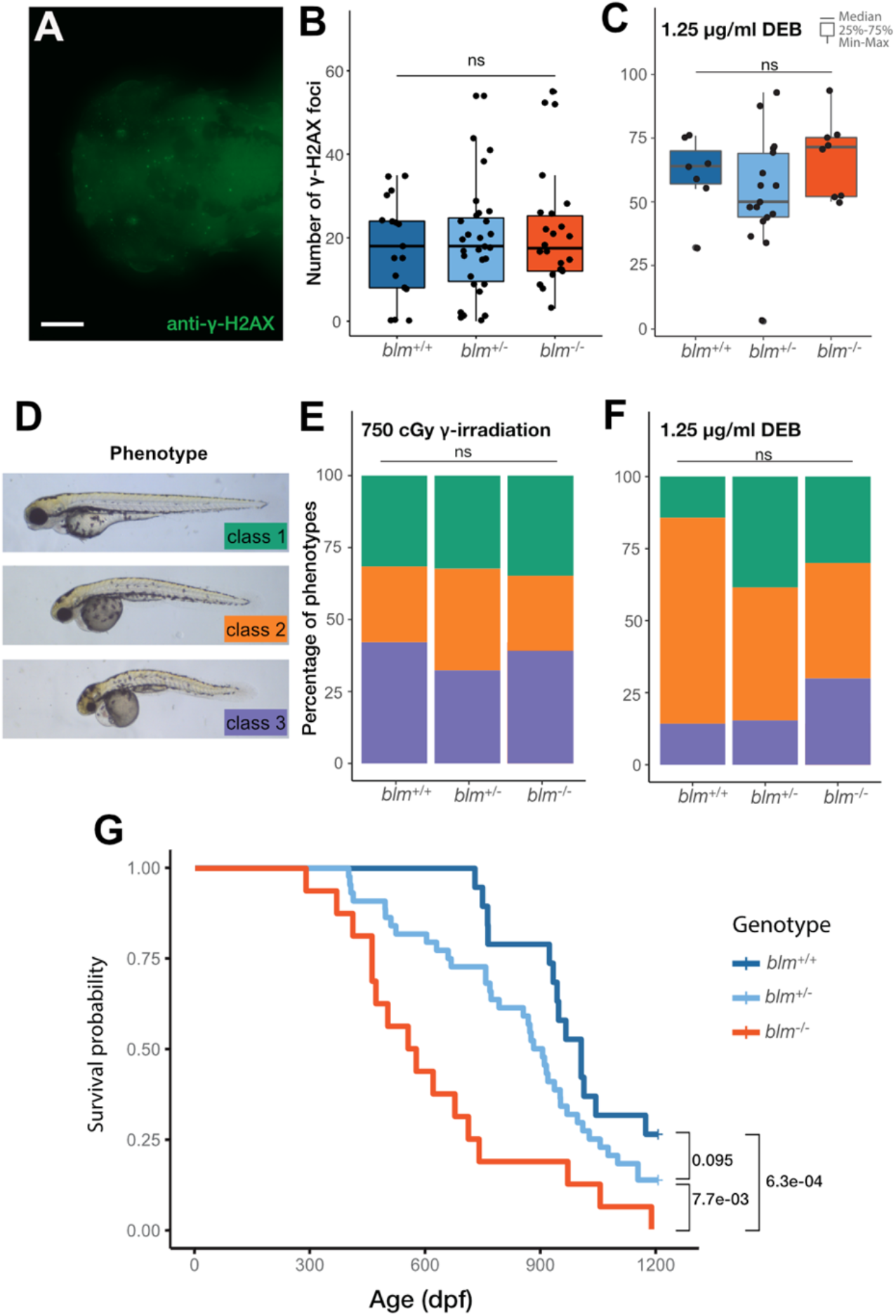
Blm loss-of-function results in increased sensitivity for DNA damage and severely reduced lifespan in zebrafish. (A) Anti-γ-H2AX labelling in the head of 2 dpf wild type embryos. (Scale bar: 100 μ m) (B,C) γ-H2AX-positive foci in the head of different *blm* genotypes under control circumstances (B) and after treatment with the DNA interstrand cross-linking agent diepoxybutane (DEB) (C). (ns – not significant). (D) Phenotypes observed after gamma irradiation: class 1 – wild type; class 2 – mild necrosis in the tectum and smaller eyes; class 3 – heavy necrosis all over the body, heart oedema. Untreated controls were all Class 1. (E-F) The distribution of phenotype severity following gamma irradiation (E) and DEB treatment (F). ns – not significant) (G) Survival probability graphs of different *blm* genotypes. (Pairwise p-values were calculated with the Log-Rank test, using Benjamini & Hochberg adjustment.)

The lack of differences in the number of DSBs prompted us to test the effects of different forms of genotoxic treatment in these animals. First, we applied 750 cGy γ-radiation on 1 dpf embryos to induce DSBs in the DNA and assayed them for phenotypic differences after one day of regeneration. Again, we observed no significant differences in the phenotypic distribution of irradiated embryos (Figure 2D,E). Next we applied diepoxybutane (DEB), a reagent with well characterized DNA-DNA and DNA-protein crosslinking effects (Ramanagoudr-Bhojappa et al., 2018). DEB treatment resulted in an overall increase in the number of γ-H2AX punctae in the heads of treated embryos (Figure 2B, C), but no discernible differences were observed between embryos of different *blm* genotypes (Figure 2C). Furthermore, while the treatment of 1 dpf embryos with 1.25 μ g/ml DEB for 24 hours resulted in phenotypes similar to those observed after γ-radiation, *blm*^−/−^ embryos showed no significant increase in the proportion of more severe phenotypes as compared to heterozygotes or wild types (Figure 2F).

Overall, we observed no short-term somatic effects of Blm deficiency under either normal or genotoxic conditions. This finding is strikingly different from the loss-of-function phenotypes of other HR-related proteins, such as Fancd1/Brca2, Fancd2, Fanci, Fancj, Fancn, Fancp, Fanct and Rad51 (Botthof et al., 2017; Ramanagoudr-Bhojappa et al., 2018).

### Blm *mutant zebrafish show markedly shortened lifespan*

In view of the fact that reduced average lifespan is one of the most striking consequences of BSyn, we set out to conduct a longevity assay on the offspring of *blm*^*+
/-*^ fish. We found that while the incidence of malignancies did not increase, *blm*^−/−^ homozygous mutant fish lived markedly shorter than wild-type animals (Figure 2G). While *blm*^+/−^ heterozygotes also appeared to have somewhat reduced lifespan compared to their wild-type siblings, this effect was not significant (Figure 2G). It has been shown in previous studies that single-allele defects of the Bloom helicase in humans lead to increased chromosome damage and elevation in the frequency of sister chromatid exchange (SCE) events. These are most likely due to the role of the protein in chromosome segregation and HR regulation, respectively (Martin et al., 1994; Salah et al., 2014). Additionally, *BLM* heterozygosity has also been linked to susceptibility to mesothelioma and breast, colorectal and endometrial cancers to differing degrees in human patients (de Voer et al., 2015; Prokofyeva et al., 2013; Schayek et al., 2017; Sokolenko et al., 2014; Sokolenko et al., 2012; Thompson et al., 2012) (Bononi et al., 2020). Furthermore, heterozygous *Blm*^+/−^ mice were reported to be more prone to develop carcinogenic abnormalities upon being subjected to oncogenic virus treatment and in tumor suppressor deficient background (Goss et al., 2002). It must be noted, however, that other human population studies found no discernible adverse effects of the *BLM*^+/−^ genotype (Cleary et al., 2003; Laitman et al., 2016).

### All blm mutant zebrafish develop into males with fertility defects

As mentioned earlier, Blm impairment did not result in compromised early viability and *blm*^−/−^ zebrafish individuals were present at expected ratios at 3 months of age. However, similar to that reported previously for most FA proteins and Rad51, complete impairment of Blm resulted in all fish developing as males, exhibiting typical morphological and behavioral (e.g. courtship) characteristics (Figure 3A). In order to reveal if the phenotype is linked to a failure in the expansion of the PGC compartment (as seen for other mentioned mutants), we crossed our *blm*^+/−^ carriers into the *Tg(ddx4:egfp)* background, where the differential expression of GFP allows for clear identification of the two sexes from ~6 weeks onwards (Krøvel and Olsen, 2002). In accordance with an all-male development, every *blm*^−/−^*;Tg(ddx4:egfp)* individual displayed little or no expression of the transgene at the age of one month, suggestive of typical male gonad development (Supplementary Figure 2).

**Figure 3:**
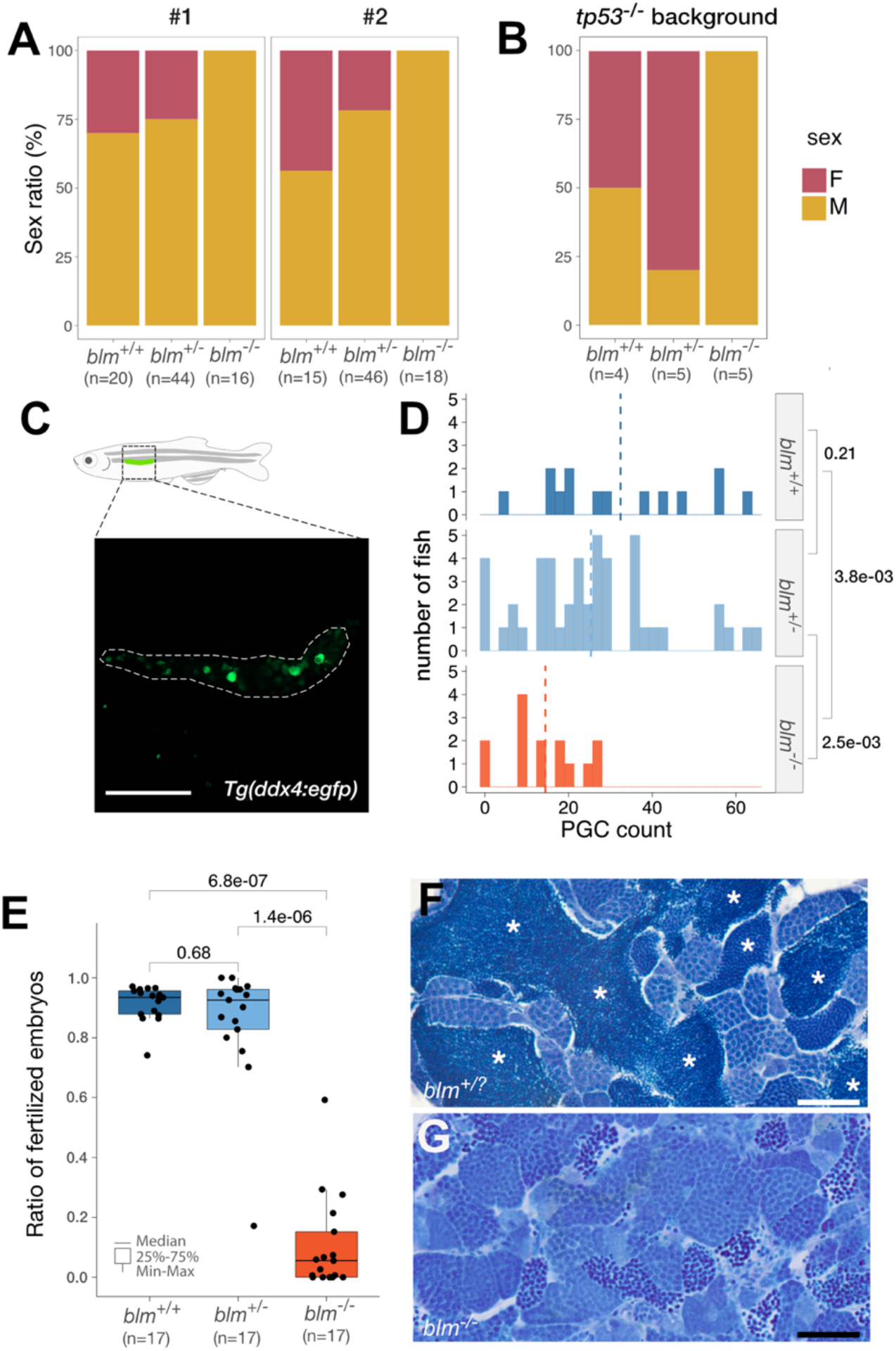
Homozygous *blm* mutants develop into males with fertility defects. (A) The sex ratio of offsprings derived from multiple *blm*^+/−^ incrosses suggests a role for Blm in zebrafish sex determination and/or gonad differentiation. (B)The male bias of *blm*^−/−^ mutant fish in the absence of a functional *tp53* indicates that the effect of Blm is Tp53-independent. (C,D) Gonadal PGC counts of different *blm* genotypes as determined on a *Tg(ddx4:egfp)* background. Juvenile fish were imaged and GFP-positive cells in the denoted area counted (C). (Scale bar: 100 μ m) Number of individuals of a given *blm* genotype (displayed on the right) showing the respective PGC counts (D). (E) The ratio of fertilized embryos in crosses using males of different *blm* genotype. (F, G) Toluidine Blue staining of testis sections from *blm*^−/−^ mutants (F) and their wild-type siblings (G). Asterisks denote spermatozoa clusters of normal densities. Mutants almost completely lacked matured spermatozoa. (Scale bar: 50 μ m).) Labels: F – female; M – male; PGC – primordial germ cell.

In order to discern whether this phenomenon was due to a precocious apoptosis of PGCs in the mutant fish, we analyzed sex ratio in *blm*^−/−^; *tp53*^*M214K*^ (Berghmans et al., 2005) double mutants and found that the male bias remained (Figure 3B). This suggests the complete masculinization of these double mutants was the result of a PGC-loss due either to Tp53-independent cell death, or to other processes not involving cell death at all. As BLM has been shown to be involved in the resolution of stalled replication forks (Manthei and Keck, 2013), and the processing of late replication intermediate structures and ultrafine anaphase-bridges (Chan et al., 2007; Sarlós et al., 2018), one possible explanation for the all-male *blm*^−/−^ phenotype is that due to the impaired cell proliferation PGC numbers in the germ lines of mutant homozygotes fail to exceed the threshold necessary for female development (Mao et al., 2010; Tzung et al., 2015a). To test this hypothesis, we counted PGCs in 2-week old *blm*^−/−^*;Tg(ddx4:egfp)* fish. In general, at this stage, in wild-types lower PGC numbers can be observed in those individuals that will eventually develop into males and higher ones in future females (Tzung et al., 2015a; Tzung et al., 2015b). Indeed we found that, in the progeny of *blm*^+/−^ carriers, homozygous mutants had significantly reduced PGC numbers compared to their wild-type and heterozygous siblings (*blm*^*+/+*^ (n=14): 32.4±18.3; *blm*^+/−^ (n=47): 25.4±16.3; *blm*^−/−^ (n=14): 14.4±9.02) (Figure 3C).

Despite the similarity between all-male phenotypes of *blm* and *fanc* mutants, we found that all *blm*^−/−^ male fish were almost completely sterile (Figure 3E), a feature that differs from that of *fanc* loss-of-function mutants (Botthof et al., 2017; Ramanagoudr-Bhojappa et al., 2018). *Blm*^−/−^ fish also displayed aberrant testis morphology (Supplementary Fig. 2) and histological analysis revealed an impairment in the formation of mature spermatozoa.

Zebrafish, similarly to other anamniotes undergo a cystic form of spermatogenesis, where single SSCs undergo mitotic clonal expansion and a final wave of meiosis to form mature spermatids (Leal et al., 2009; Yoshida, 2016). While in *blm*^−/−^ males cysts of different size comprising differing number of spermatogonia were present, post-meiotic spermatid clusters were extremely rarely observed (Figure 3F,G).

Our results, therefore, suggest a failure of spermatogonial cells to complete meiosis in the absence of Blm function.

### Blm-deficient spermatids are stuck in meiotic prophase I

Blm has been implicated in multiple steps of the meiotic process (Walpita et al., 1999). Therefore, we sought to understand whether the lack of spermatids in the testis of *blm*^−/−^ males is due to defective meiosis. Labeling DSBs with anti-γ-H2AX suggests that unlike in *spo11* mutants (Blokhina et al., 2019), in *blm*^−/−^ testes DSB formation is not affected (Supplementary Fig. 3). We examined meiosis using an antibody specific for Sycp3, a component of the synaptonemal complex, previously shown to accumulate during the meiotic prophase (Blokhina et al., 2019). While in the testes of wild-type fish spermatids showed a typical mix of Sycp3 staining patterns, in those of *blm*^−/−^ mutant individuals nuclei either displayed Sycp3 patterns typical for leptotene, or showed an aberrant accumulation of Sycp3 in the nuclear periphery (Figure 4A-C). The latter pattern has been observed before in apoptotic spermatids of rats (Escobar et al., 2019), suggesting that in *blm*^−/−^ mutants spermatogonia are stuck in early meiotic phases and undergo cell death. Supportive of this, in *blm*^−/−^ mutants spermatocytes in spermatogenic cysts often undergo programmed cell death as visualized by Caspase-3 staining (Figure 4D-F). This phenotype could be attributed to the well-documented loss-of-function of BLM in the dissolution and/or resolution of double Holliday junctions, late intermediates of HR (Agostinho et al., 2013; O’Neil et al., 2013; Schvarzstein et al., 2014; Wu and Hickson, 2003). Accordingly, in the absence of the yeast BLM ortholog, Sgs1, the number of joint molecules made of up to four interconnected duplexes is increased (Oh et al., 2007), while in mice the lack of Blm results in excess chiasmata and mispairing between nonhomologous chromosomes in spermatocytes (Holloway et al., 2010).

**Figure 4:**
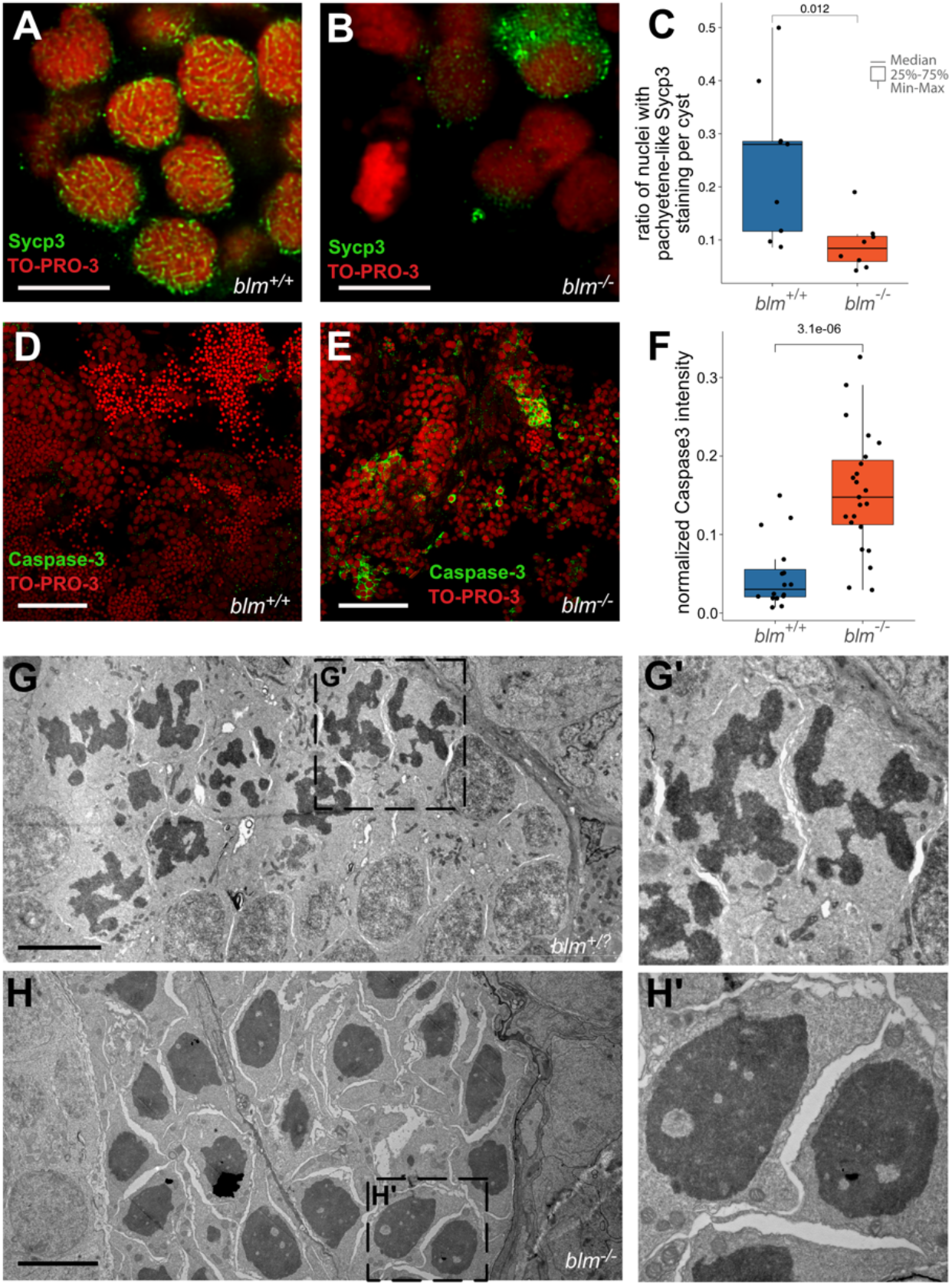
Blm loss-of-function results in meiotic defects during spermatogenesis in zebrafish males. (A,B) Representative Sycp3 immunostainings of wild type and *blm*^−/−^ spermatogonial cysts. In wild type cysts (A) patterns typical for meiotic prophase I can be detected, whereas in mutant cysts (B) aberrant Sycp3 patterns (green) can be observed. TO-PRO-3 staining (red) denotes nuclei. (Scale bar: 8 μ m). (C) Ratio of nuclei showing Sycp3 immunostaining patterns characteristic for pachytene. Comparison of wild type and mutant testes suggests that *blm*^−/−^ are defective in entering pachytene. (D, E) Cells undergoing programmed cell death as shown by Caspase-3 (green) staining in wild type (D) and *blm*^−/−^ testes (E). TO-PRO-3 staining (red) denotes nuclei. (Scale bar: 50 μ m) (F) Normalized Caspase-3 stainings of wild type and mutant testes suggest increased apoptosis in the absence of Blm. (G, G’) Electron microscopic images of representative spermatogenic cysts in wild-type males. (Scale bar: 5 μ m) (H, H’) Electron microscopic images of representative spermatogenic cysts in *blm*^−/−^ mutant males showing an aberrant morphology. (Scale bar: 5 μ m)

To further assess possible defects in the segregation of meiotic chromosomes, we performed electron microscopic (EM) imaging of germ cells. While in single spermatogenic cysts of wild-type and heterozygote testes we observed spermatocytes at different phases of cell cycle (Figure 4G,G’), often with well-delineated, clearly separated chromosomal structures, in *blm*^−/−^ mutants an aberrant chromatin “blob” appeared in most spermatids, suggesting that their chromosomes failed to segregate and remained entangled (Figure 4H,H’) (Hatkevich et al., 2017).

### Discussion

The effects of BSyn on human lifespan and fertility are well documented. Here we describe and validate a novel zebrafish model for this rare disease. Our new model recapitulates several features of human BSyn (Adam et al., 1993; Cunniff et al., 2017), including shortened lifespan and infertility. This new line also provides important insights into the pathomechanism of the meiotic defects associated with Blm loss-of-function and expands our understanding of the formation and differentiation of zebrafish gonads.

There are several explanations why mutations in *BLM* could affect human lifespan. Accumulation of mutations due to an inability to repair double-stranded DNA breaks that arise as a result of environmental genotoxic effects is one plausible explanation. Indeed, mutations of multiple *fanc* genes and *rad51*, all involved in DSB repair, result in an increase of cell death in zebrafish upon exposure to genotoxic agents (Botthof et al., 2017; Ramanagoudr-Bhojappa et al., 2018). Yet, our results show that genotoxic treatments do not result in more DNA lesions or more severe phenotypes in *blm*^−/−^ mutant zebrafish (Figure 2A-F). Compared to their wild-type and heterozygote siblings, however, homozygous *blm*^−/−^ zebrafish have a significantly shorter lifespan (Figure 2G).

The function of RecQ helicases during HR events is well documented. In bacteria RecQ is required to reduce the number of illegitimate recombination events (Ferencziová et al., 2018; Hanada et al., 1997; Harami et al., 2017) and in mammals BLM is involved in the dissolution of double Holliday junctions during HR, and the processing of late replication intermediates, especially at fragile sites (Bizard and Hickson, 2014; Oh et al., 2007; Wu and Hickson, 2003). The impairment in the resolution of DNA junctions might contribute to an elevated mutation rate that results both in the shortened lifespan and an increase in the number of malignancies, both in human and mice (Chan et al., 2018; Goss et al., 2002; Gruber et al., 2002). Interestingly, we did not observe an increase in the incidence of cutaneous tumors in *blm*^−/−^ mutants and *blm*^+/−^ carriers during our lifespan analysis (not shown), although occasional testicular tumors have been observed amongst the *blm*^−/−^ males. While it is possible that due to the nature of our protocol we were not able to account for an increase in internal (e.g. intestinal) tumors, our results also suggest that impairment of Blm function has other effects than just a simple increase in the mutation rate.

Our results clearly show the importance of Blm in the context of somatic tissues but even more interestingly we were also able to demonstrate multiple roles for Blm in zebrafish gametogenesis and SD/SDiff (Figure 5), as *blm*^−/−^ individuals invariably develop into males with severely compromised fertility (Figure 3A,E). We provide evidence that this is due to the roles of Blm in both mitotic and meiotic events.

**Figure 5:**
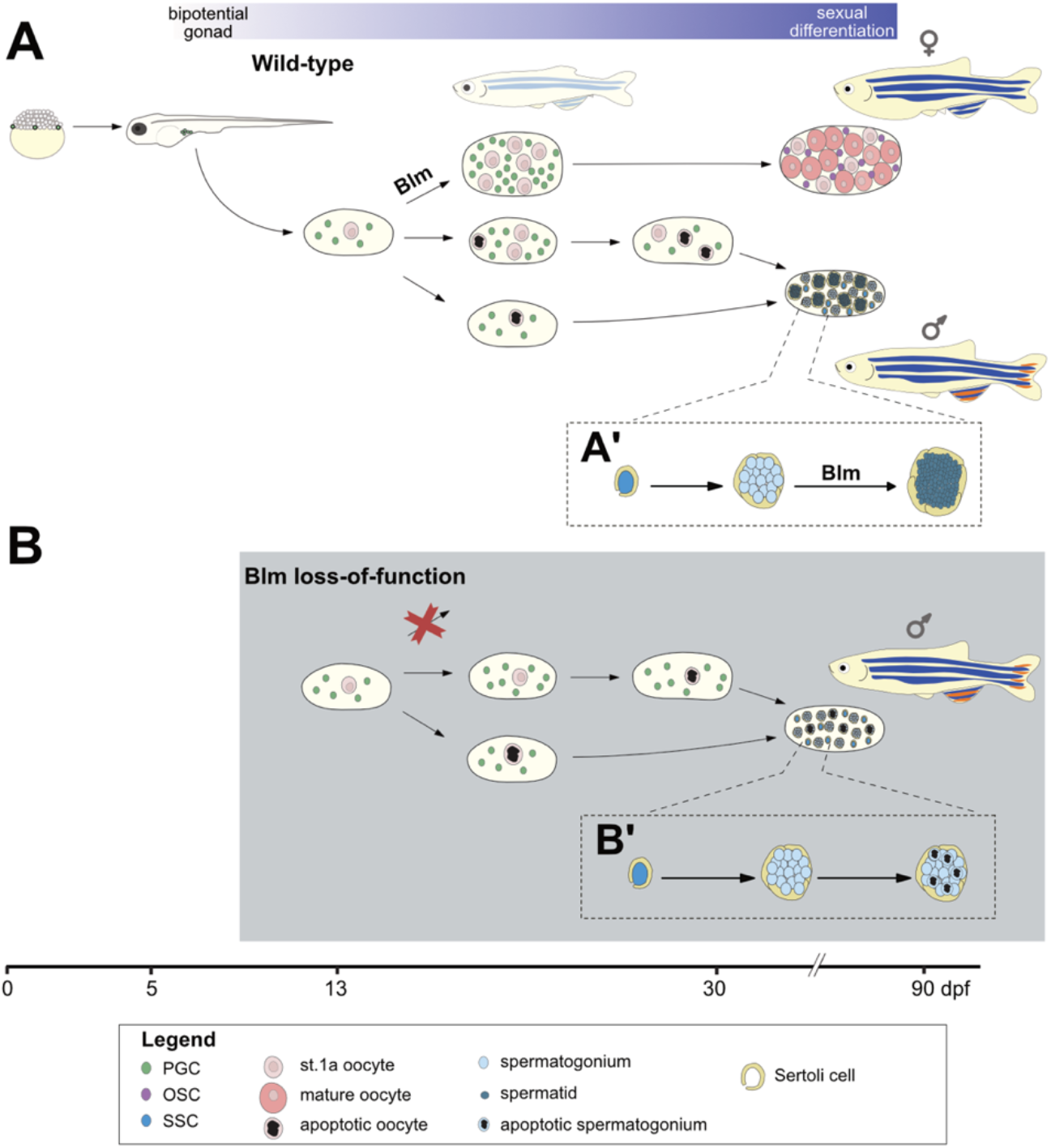
A model for the multiple roles of Blm during zebrafish gonadal development. During the early stages of ontogenesis *vasa* expressing primordial germ cells (PGC) proliferate and migrate from their initial positions adjacent to the yolk syncytial layer to the gonadal mesoderm. There their proliferation continues, while a varying number differentiate into stage 1a oocytes resulting in the development of the ‘juvenile ovary’. (A) In wild-type fish the number of PGCs may further increase to the point where it crosses the threshold required for female fate determining pathways to activate, resulting in the development of mature ovaries with PGC-derived ovarian stem cells (OSC) and mature oocytes. If the threshold is not reached either due to insufficient PGC expansion, or none at all, a major shift in gonadal environment occurs through which oocytes (and possibly even a subset PGCs) are eliminated via apoptosis. Subsequently, the formation of spermatogonial stem cells (SSC) and Sertoli cells occurs either by (i) differentiation of common OSC and SSC precursors (gonadal stem cells) into SSCs that in turn induce Sertoli cell differentiation in the soma; or (ii) by the transdifferentiation of somatic follicular cells into Sertoli cells that promote SSC differentiation and expansion from gonadal stem cells, thus leading to testis development. (A’) In the wild-type testis SSCs are enveloped by Sertoli cells where they further differentiate into spermatogonia. In this structure, known as a cyst, spermatogonia go through mitotic expansion followed by synchronous meiosis into spermatids. (B) In Blm loss-of-function individuals the number of PGCs never reaches the threshold necessary for female development, therefore all homozygous mutants develop as males. (B’) An explanation for the subfertility of *blm*^−/−^ males may be that in the absence of Blm accurate DNA repair during spermatogonial meiosis is hindered, and therefore cell death is induced.

In zebrafish one of the most important elements of SD/SDiff is the number of PGCs in the developing gonads. If the PGC population fails to undergo a mitotic expansion during metamorphosis, the juvenile bipotential ovary almost invariably develops into a testis (Tzung et al., 2015a; Tzung et al., 2015b; Ye et al., 2019). Replication stress that naturally arises in some dividing PGCs during mitosis normally would activate DNA repair pathways, and the impairment of these pathways leads to a failure in PGC expansion (Luo et al., 2014). This explains why the mutations of genes that control the major DNA repair pathways, such as those that encode the components of the FA complex or Rad51 (Figure 5A), all result in a sex-reversal phenotype in zebrafish (Botthof et al., 2017; Ramanagoudr-Bhojappa et al., 2018; Rodríguez-Marí et al., 2011; Shive et al., 2010). (For the list of those genes whose loss-of-function results in male bias in zebrafish see Supplementary Table S1.)

One important difference, however, between the sex-reversal observed in *blm*^−/−^ zebrafish compared to that of *fancd1/brca2, fancj* and *rad51* mutants (Botthof et al., 2017; Ramanagoudr-Bhojappa et al., 2018; Rodríguez-Marí et al., 2011; Shive et al., 2010) is that the all-male phenotype for the latter is dependent on Tp53 function. Crossing these mutants into a *tp53* mutant background was sufficient to rescue sex-reversal (through the rescue of PGC numbers), however, all *blm*^−/−^*;tp53*^−/−^ double mutants developed as males. We hypothesize that in the absence of Blm DNA damage can accumulate in some fast proliferating PGCs leading to mitotic arrest and spindle checkpoint activation, triggering mitotic cell death. For unknown reasons, however, Blm loss-of-function does not appear to trigger Tp53 activation and it might lead to apoptosis through a Tp53-independent mechanism, probably through the activation of one of the Tp53-homologs, for example TAp73 (Toh et al., 2010).

Similarly to Blm, some factors are not only involved in mitotic DNA repair, but also have important roles in regulating homologous recombination during meiosis (Hunter, 2015). The impairment of these proteins, such as Fancd1/Brca2, Fancj and Rad51 leads to defects in both cell division types. Thus, similarly to *blm*^−/−^ mutants, both sex reversal and male subfertility can be observed in the mutants of the genes that encode these factors (Botthof et al., 2017; Ramanagoudr-Bhojappa et al., 2018; Rodríguez-Marí et al., 2011; Shive et al., 2010). In contrast, only subfertility was observed in the mutants of *hsf5, mlh1* and *spo11* all of which are involved in the meiotic process (Blokhina et al., 2019; Feitsma et al., 2007; Saju et al., 2018) (Figure 5A).

The role of Blm in meiosis is well established, hence it is not entirely unexpected that we observe defects during spermatogenesis in zebrafish *blm*^−/−^ mutants. In *C. elegans* BLM (HIM-6) is required to convert licensed cross-over (CO) designated sites into *bona fide* COs and in *him-6* mutants abnormal chromosomal segregation can be observed (Agostinho et al., 2013; O’Neil et al., 2013; Schvarzstein et al., 2014). Similarly, loss of *DmBlm* results in excessive number of COs, nondisjunction and aneuploidy during *Drosophila* meiosis as well (Hatkevich et al., 2017). In mice *Blm* expression is upregulated during the leptotene stage (Chen et al., 2018) and its impairment leads to defects in meiotic progression due to the improper pairing and synapsis of homologous chromosomes and BLM-deficient cells often undergo apoptosis during spermatogenesis (Holloway et al., 2010). This latter phenotype is highly reminiscent to what we observe in the testes of our *blm*^−/−^ mutant zebrafish (Figure 4).

Fertility problems have been also reported for BSyn patients of both sexes. Interestingly, the phenotype seems to be milder in women, where delayed puberty and early menopausal arrest could contribute to the observed subfertility, but multiple cases of affected women delivering healthy offspring have been described in the literature (Adam et al., 1993; Cunniff et al., 2017). In contrast, males are invariably infertile, with problems in pubertal progression and severe oligospermia or azoospermia (Adam et al., 1993; Cunniff et al., 2017).

There are a number of outstanding questions. The viability and relatively mild somatic phenotype of Blm loss-of-function suggest that there is redundancy with other RecQ orthologs during the division of somatic cell progenitors. Based on their expression profiles (Figure 1B) *recql* and *recql5* could have important roles during embryogenesis, but it remains to be determined how embryonic and larval development (and viability) are affected when these genes are mutated. Also, while the expression of *blm* is clearly elevated in zebrafish oocytes, *recql4* has a similar expression pattern (Figure 1B) (White et al., 2017; Winata et al., 2018). It also remains to be seen to what extent are their functions redundant during gametogenesis.

Finally, it remains to be determined if the role of Blm in SD/SDiff could be related to the genomic peculiarities of the zebrafish SD system. Notably, although domesticated strains lost their heterogametic sex chromosome (W), a locus on Chromosome 4 linked previously to SD in wild animals was shown to be likely involved in the SD of laboratory fish as well (Ortega-Recalde et al., 2019; Wilson et al., 2014). This region contains a maternal specific rDNA cluster, dubbed fem-rDNA, exhibiting female-specific total demethylation and amplification during oocyte development (Ortega-Recalde et al., 2019). The fem-rDNA region is rich in guanine quadruplex (G4) regions and is flanked by immunoglobulin switch-like recombination regions (Breit et al., 2020). As Blm has been implicated in the replication of repetitive regions such as rDNA clusters (Schawalder et al., 2003), in rDNA maintenance and transcription (Tangeman et al., 2016), as well as in the unwinding of G4-containing DNA regions (Chatterjee et al., 2014; van Wietmarschen et al., 2018; Wu et al., 2015), it is possible that *blm*^−/−^ homozygous mutants are unable to amplify this cluster and this could affect the early mitotic amplification of the PGCs.

In summary we describe here a novel zebrafish model for BSyn which recapitulates several important features (e.g. sterility and reduced lifespan) of the human condition and also provides important insights into the biological function of Blm. Surprisingly, we also show that lack of Blm function does not increase the number of DSB events under genotoxic conditions. As Blm also affects zebrafish SD/SDiff through the regulation of PGC number, our results reveal that both meiosis and mitosis can be affected in the absence of this helicase.

## Materials and methods

### Zebrafish husbandry

Wild-type (AB), and *blm*^*elu10*^ and *tp53*^*M214K*^ (Berghmans et al., 2005) mutant zebrafish in non-transgenic and *Tg(ddx4:egfp)* (Krøvel and Olsen, 2002) backgrounds used in this study were maintained and bred in the fish facility of ELTE Eötvös Loránd University according to standard protocols (Aleström et al., 2019; Westerfield, 2000). All experimental procedures were approved by the Hungarian National Food Chain Safety Office (Permit Numbers: PE/EA/290-2/2018, PE/EA/2025-7/2017). Fish involved in the lifespan assay were reared at similar stocking densities.

### CRISPR/Cas9 mutagenesis and genotyping

SgRNA targeting the sequence 5’-AAGACAAATACAGTTAATCC-3’ in the 4^th^ exon of *blm* was synthesized and injected as described before (Gagnon et al., 2014). In the P0 generation we isolated mosaic fish that carried in their germline the c.302_304delACAAinsTT indel mutation. These carriers were outbred with AB wild-type individuals. Genomic DNA isolation for genotyping of fin clips from adults was carried out as described before (Meeker et al., 2007). Genotyping for the *blm*^*elu10*^ allele was accomplished via allele-specific PCR. Two reactions were assembled per sample, one with wild-type specific (5’-CGCCCAGCAAACCGAAGACAA-3’) and another with mutant specific (5’-GCGCCCAGCAAACCGAAGTTA-3’) forward primers. In both cases the reverse primer was: 5’-GTATCCGGTGAAACACAGTGGA-3’. Genotypes were determined from gel electrophoretic analysis. Genotyping of the *tp53*^*M214K*^ allele was performed by Sanger sequencing the PCR product created with the following forward and reverse primers: 5’-TTGCCAGAGTATGTGTCTGTCC-3’ and 5’-CAGCATCATGAAGCATCAAA-3’, respectively.

### Genotoxic treatments

γ-irradiation of fish was carried out as described before (Botthof et al., 2017) at the Basic Medical Science Center of Semmelweis University, Budapest. DEB treatments were performed as described before (Ramanagoudr-Bhojappa et al., 2018).

### Sexing zebrafish

The sex of individual zebrafish was determined first based on sexually dimorphic phenotypic traits in adults and confirmed later by visual inspection and/or histological analysis of their dissected gonads.

### Histology

For histological examination, fish were dissected and their testes were fixed by fixative containing 4 % paraformaldehyde. The samples were embedded using JB-4 resin (Merck-Sigma, EM0100) as described in the manufacturer’s protocol. 2–5 μm thin sections were made by rotation microtome and stained with Toluidine Blue. (Briefly, 1% Azure B, and 1% Toluidine Blue, 1% borax stock solutions were mixed together and saccharose added to a final 0.5% concentration. Sections were heated and stained for 20-30 minutes.) Sections were examined and photographed using a Zeiss Axio Imager microscope system.

For immunostaining we used the anti-Sycp3-rabbit (Abcam, ab15093; 1:500 dilution), anti-Caspase3-rabbit (Cell Signaling Technology, 9661l; 1:500 dilution) and anti-γH2AX-rabbit (GeneTex, GTX127342; 1: 1000 dilution in larvae, 1:200 dilution in testes) primary antibodies in combination with anti-rabbit-Alexa488 (Invitrogen, A-11008; 1:200 dilution) secondary antibody. Nuclei were stained with TO-PRO-3 (Topro, Invitrogen, T3605; 1:3000 dilution).

Larvae stained with anti-γH2AX were imaged on a Zeiss AxioImager M2 microscope, with a Zeiss EC Plan-NEOFLUAR 10X / 0.3 NA objective and a Zeiss Colibri 7 light source.

In the case of the anti-γH2Ax and anti-Caspase3 immunostainings statistical analysis has been made based on approximately 20 nonoverlapping pictures with the same size. Using ImageJ software, we measured the integrated pixel density of every record for the 488 nm channel and normalized these values by the integrated pixel density value of the Topro (661 nm) channel.

For electron micrography we fixed the samples using a solution containing 3.2% PFA, 0.2% glutaraldehyde, 1% sucrose, 40 mM CaCl2 in 0.1 M cacodylate buffer. Post-fixation was performed with 1% ferrocyanide-reduced osmium (White et al., 1979). Following dehydration by graded ethanol series, the samples were embedded into Spurr low viscosity epoxy resin medium. Ultrathin sections were made and collected on Formvar coated grids. Sections were then counterstained with 2% uranyl acetate solution and Reynolds’s lead citrate solution. For examination, a JEOL JEM 1011 transmission electron microscope was used, equipped with a Morada 11-megapixel camera.

*In situ* hybridizations were performed has been described before (Bellipanni et al., 2006). A ~1 kb fragment of the *blm* cDNA was cloned using primers blm-F – 5’-GGAGTCGAAACACCTGGTGGTA-3’ and blm-R – 5’-CTCATCAATGACCAAGCGAGCC-3’ into pGEM-T vector (Promega) following the manufacturer’s protocol, cut with SacII and transcribed using SP6 polymerase.

### PGC quantification

The offspring of *blm*^+/−^*;Tg(ddx4:egfp)* fish were raised to 5-6 mm standard length and were fixed in 4 % PFA in PBS. Larvae were screened for *GFP* expression under a Zeiss SteREO Lumar.V12 microscope equipped with a NeoLumar S 0.8x objective and a CoolLED pE-300lite light source. GFP-positive specimens were embedded in 2 % low melting point agarose gel and were imaged under a Zeiss Axio Imager M2 upright microscope with semi-confocal ApoTome 2.0 using an EC Plan-NeoFluar 10x 0.3 objective and a Colibri 7 LED light source. Standard length was measured following imaging procedures. PGCs were manually enumerated in Zeiss ZEN.

### Fertility assay

Male fish for all *blm* genotypes were individually bred with wild-type *AB* females of similar age (8 months post fertilization). Fish were paired at random in repeated attempts with at least 5 days of rest between breeding sessions. Collected eggs were analyzed at least 6 hours post-fertilization, based on the occurrence of epiboly. Decomposing eggs were excluded from statistical analysis, as success of fertilization could not be determined.

### Statistics and visualization

Multiple sequence alignments were performed using the Uniprot UGENE toolkit (Okonechnikov et al., 2012). For the alignments we used sequences for the helicase ATPase and helicase C-terminal domains of the respective proteins. Statistical analysis and visualization was performed in R (R Core Team, 2018) using the *ggplot2* package (Wickham, 2016). To display phylogenetic relationships we used the *jsPhyloSVG* script (Smits and Ouverney, 2010). For lifespan analysis we used the *survminer* package (v. 0.4.6, Kassambara et al., 2019). All figures have been assembled in Affinity Designer (Serif Europe).

## Funding

This work was funded by grant NKFIH K-116072 from the Hungarian National Research, Development and Innovation Office (to M.K. and M.V.). G.M.H. is supported by the Premium Postdoctoral Program of the Hungarian Academy of Sciences (Grant PREMIUM-2017-17). The research project was part of the ELTE Thematic Excellence Programme 2020 supported by the National Research, Development and Innovation Office (TKP2020-IKA-05). LO was supported by the Frontline Research Excellence Grant of the National Research, Development and Innovation Office of Hungary (KKP 126764).

## Acknowledgements

We thank Anita Rácz and other members of the Fish Genetics Group at ELTE for fish care and valuable discussions of the results. We also thank István Katona for his gift of the anti-Caspase3 antibody and Lisbeth Olsen for the permission to use the *Tg(ddx4:egfp)* line in our study. Finally, we thank Balázs Bényei at the Semmelweis University Basic Medical Science Center for his help with the irradiations.

## Author contributions

Conceptualization: Mihály Kovács, Gábor Harami, László Orbán, Máté Varga. Data curation: Máté Varga, Tamás Annus, Bálint Jezsó. Funding acquisition: Mihály Kovács, Máté Varga, László Orbán.Investigation: Tamás Annus, Dalma Müller, Bálint Jezsó, György Ullaga, Gábor Harami, Máté Varga. Methodology: Máté Varga, Tamás Annus, Bálint Jezsó, László Orbán. Project administration: Mihály Kovács, Máté Varga. Supervision: Mihály Kovács, Máté Varga. Writing – original draft: Tamás Annus, Mihály Kovács, László Orbán, Máté Varga.

## SUPPLEMENTARY INFORMATION

**Supplementary Figure S1:**
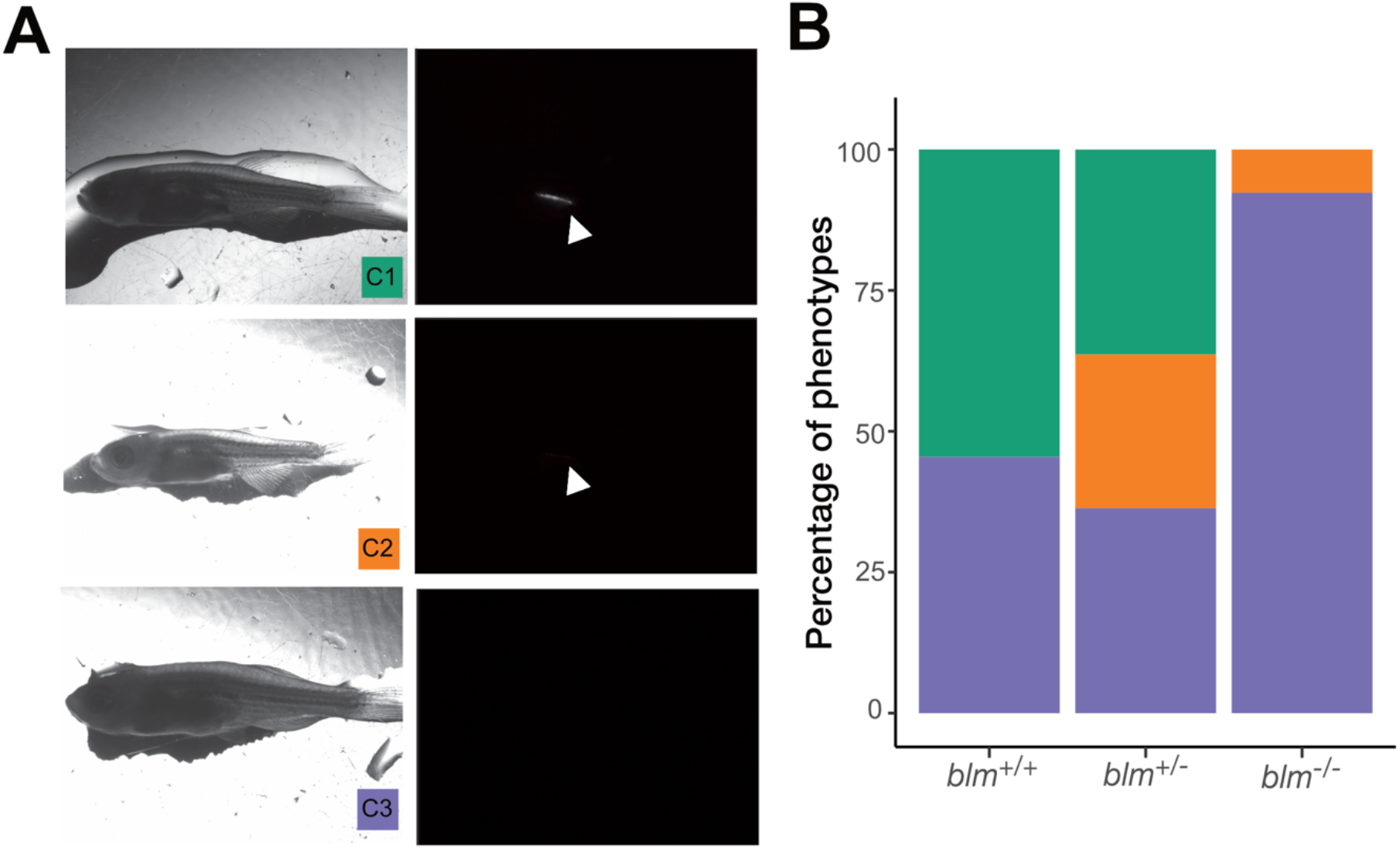
In the absence of Blm, PGCs do not expand. (A) At one month of age the gonads of *Tg(ddx4:egfp)* zebrafish show either strong fluorescence (C1), weak fluorescence (C2) or no fluorescence (C3). (B) While in *blm*^*+/+*^*;Tg(ddx4:egfp)* and *blm*^+/−^*;Tg(ddx4:egfp)* zebrafish the number of strong expressing fish was comparable to those of not expressing the transgene, gonads of *blm*^−/−^*;Tg(ddx4:egfp)* fish almost never showed fluorescence.

**Supplementary Figure S2:**
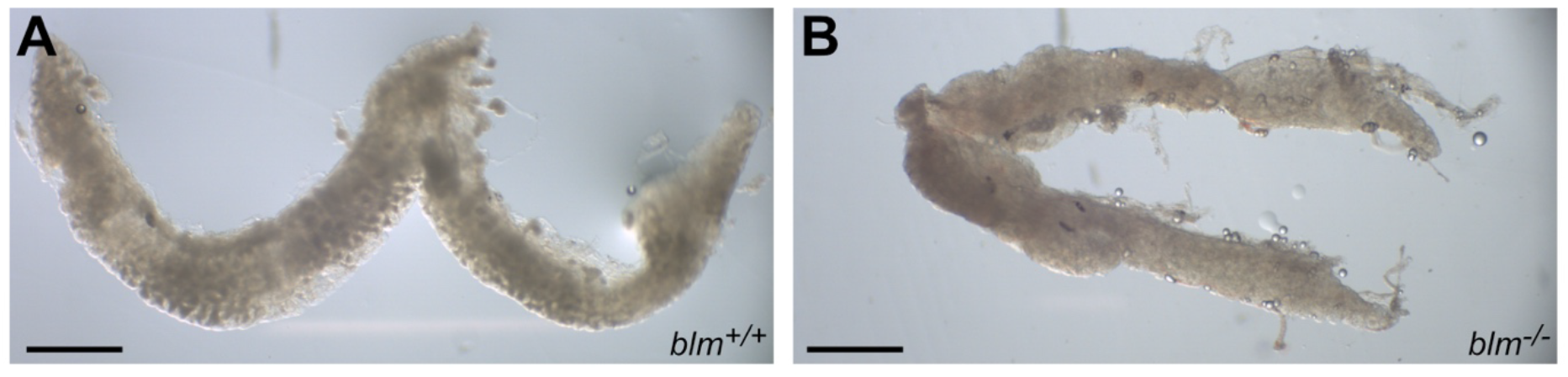
Testes of *blm* mutants show abnormal morphology. Compared to wild type testes (A) those of *blm*^−/−^ individuals (B) show less pronounced seminiferous tubule structure under transmitted light stereo microscopy and in some cases are significantly shorter. (Scale bar: 1 mm)

**Supplementary Figure S3:**
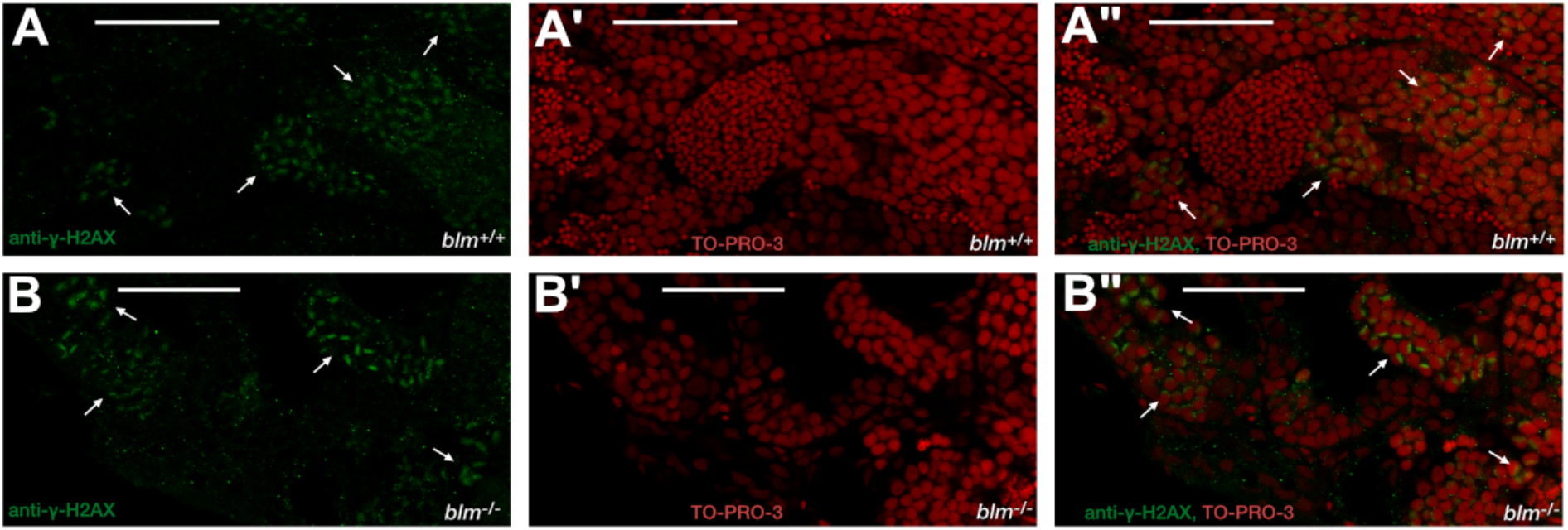
DSBs occur in meiotic spermatogonia of *blm* mutants. (A) (A-A”) Anti-γ-H2AX and nuclear (TO-PRO-3) labelling of wild type testes shows the presence of DSBs in spermatogonia that enter meiotic prophase I. (White arrows show clusters of spermatogonia with DSBs.) (Scale bar: 50 μ m) (B-B”) Anti-γ-H2AX and nuclear (TO-PRO-3) labelling of *blm* mutant testes suggest that DSB formation is impaired in these mutants. (White arrows show clusters of spermatogonia wit DSBs.) (Scale bar: 50 μ m)

**Supplementary Table S1:**
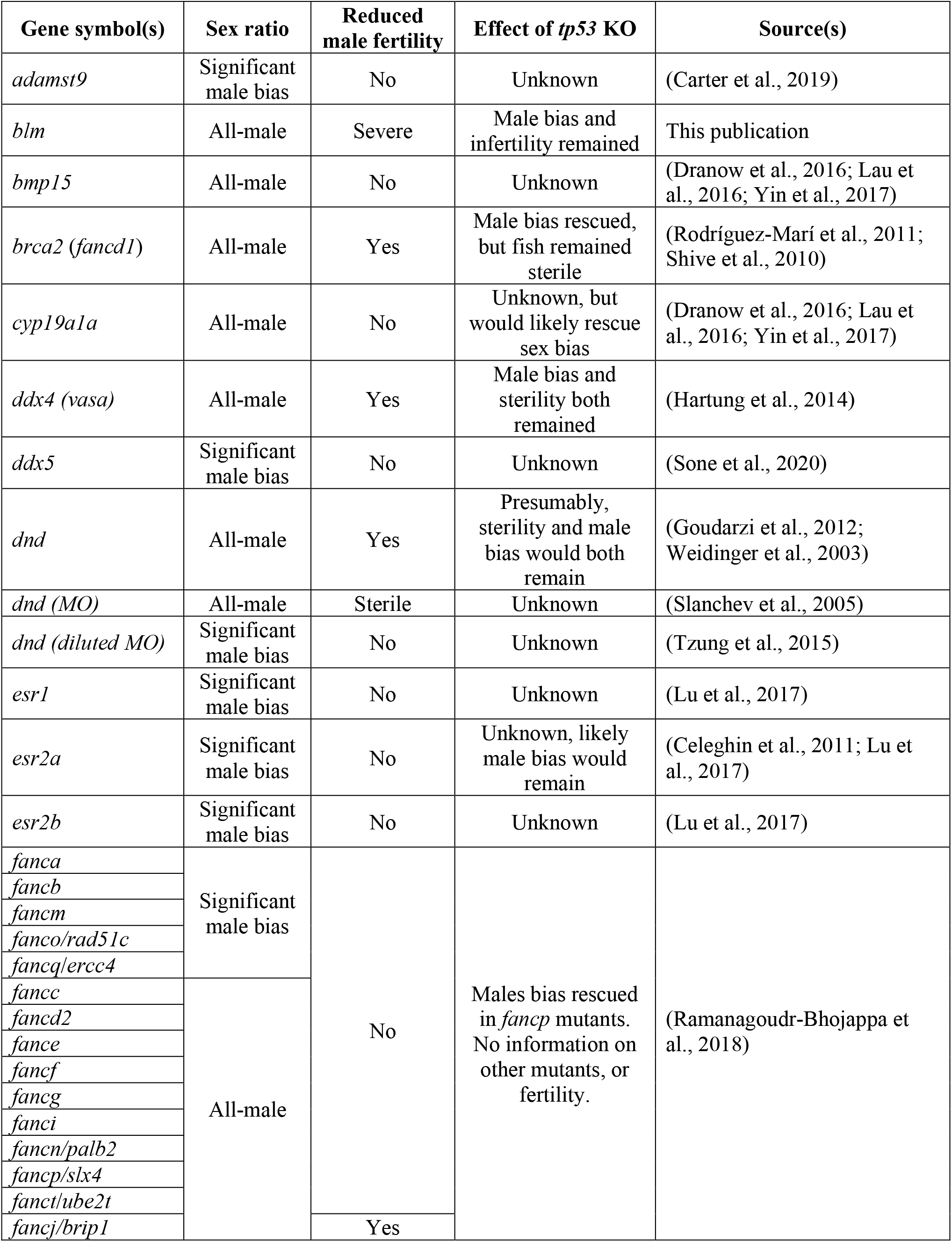
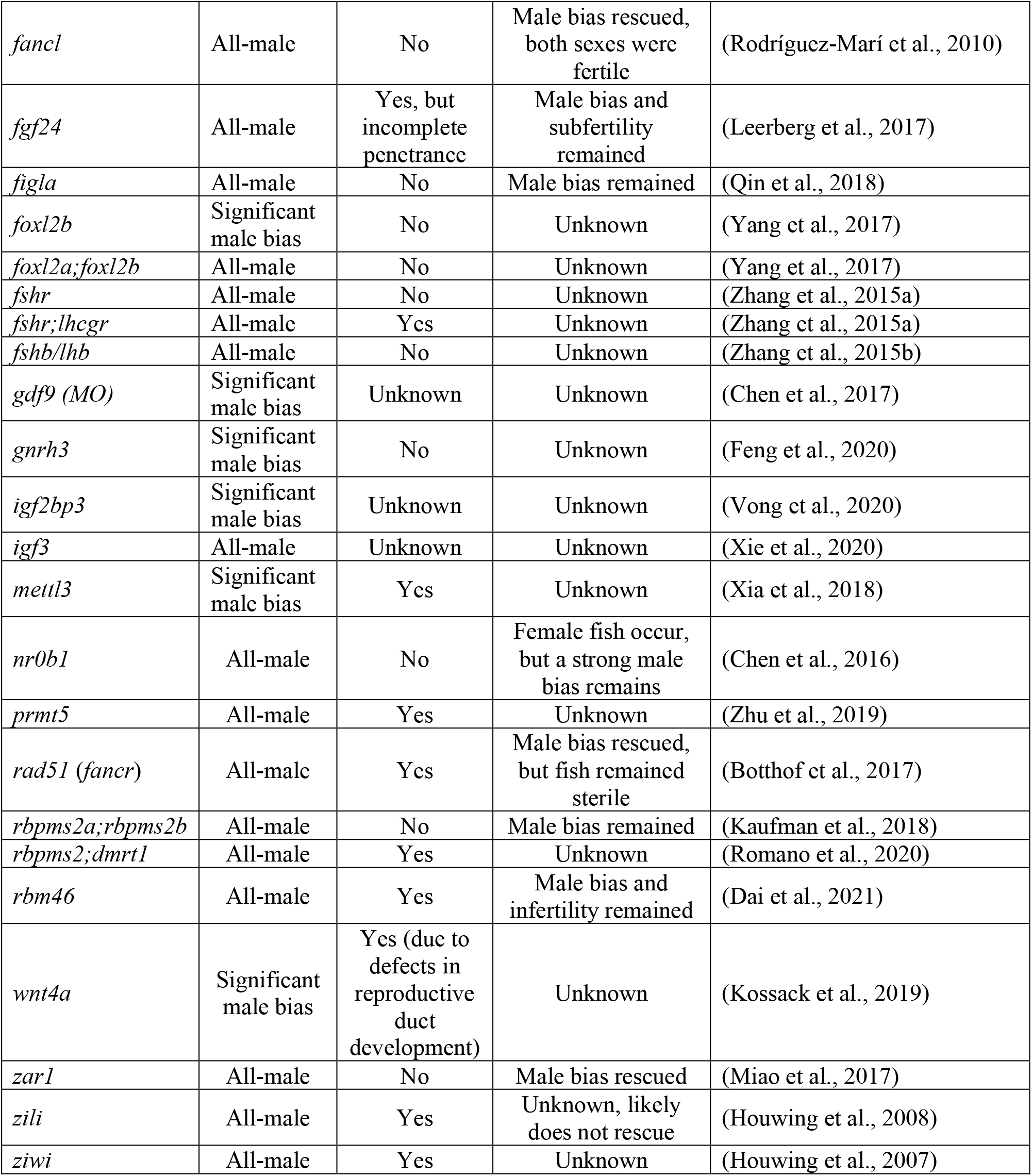
Loss-of-function conditions causing male bias in zebrafish.

## References

Adam, M. P., Ardinger, H. H., Pagon, R. A., Wallace, S. E., Bean, L. J., Stephens, K., Amemiya, A., Flanagan, M. and Cunniff, C. M. (1993). Bloom Syndrome. GeneReviews® [Internet] Seattle (WA): University of Washington, Seattle; 1993–2021.

Agostinho, A., Meier, B., Sonneville, R., Jagut, M., Woglar, A., Blow, J., Jantsch, V. and Gartner, A. (2013). Combinatorial regulation of meiotic holliday junction resolution in C. elegans by HIM-6 (BLM) helicase, SLX-4, and the SLX-1, MUS-81 and XPF-1 nucleases. PLoS Genet 9, e1003591.

Aleström, P., D’Angelo, L., Midtlyng, P. J., Schorderet, D. F., Schulte-Merker, S., Sohm, F. and Warner, S. (2019). Zebrafish: Housing and husbandry recommendations. Lab. Anim. 51, 23677219869037.

Anderson, J. L., Rodríguez-Marí, A., Braasch, I., Amores, A., Hohenlohe, P., Batzel, P. and Postlethwait, J. H. (2012). Multiple sex-associated regions and a putative sex chromosome in zebrafish revealed by RAD mapping and population genomics. PLoS ONE 7, e40701.

Aubrey, B. J., Kelly, G. L., Janic, A., Herold, M. J. and Strasser, A. (2018). How does p53 induce apoptosis and how does this relate to p53-mediated tumour suppression? Cell Death Differ 25, 104– 113.

Bellipanni, G., Varga, M., Maegawa, S., Imai, Y., Kelly, C., Myers, A. P., Chu, F., Talbot, W. S. and Weinberg, E. S. (2006). Essential and opposing roles of zebrafish beta-catenins in the formation of dorsal axial structures and neurectoderm. Development 133, 1299–1309.

Berghmans, S., Murphey, R. D., Wienholds, E., Neuberg, D., Kutok, J. L., Fletcher, C. D. M., Morris, J. P., Liu, T. X., Schulte-Merker, S., Kanki, J. P., et al. (2005). tp53 mutant zebrafish develop malignant peripheral nerve sheath tumors. Proceedings of the National Academy of Sciences 102, 407–412.

Bizard, A. H. and Hickson, I. D. (2014). The dissolution of double Holliday junctions. Cold Spring Harb Perspect Biol 6, a016477–a016477.

Blokhina, Y. P., Nguyen, A. D., Draper, B. W. and Burgess, S. M. (2019). The telomere bouquet is a hub where meiotic double-strand breaks, synapsis, and stable homolog juxtaposition are coordinated in the zebrafish, Danio rerio. PLoS Genet 15, e1007730.

Bloom, D. (1954). Congenital telangiectatic erythema resembling lupus erythematosus in dwarfs; probably a syndrome entity. AMA Am J Dis Child 88, 754–758.

Bononi, A., Goto, K., Ak, G., Yoshikawa, Y., Emi, M., Pastorino, S., Carparelli, L., Ferro, A., Nasu, M., Kim, J.-H., et al. (2020). Heterozygous germline BLM mutations increase susceptibility to asbestos and mesothelioma. Proc Natl Acad Sci USA 117, 33466–33473.

Botthof, J. G., Bielczyk-Maczyńska, E., Ferreira, L. and Cvejic, A. (2017). Loss of the homologous recombination gene rad51 leads to Fanconi anemia-like symptoms in zebrafish. Proc Natl Acad Sci USA 114, E4452–E4461.

Breit, T. M., Rauwerda, H., Pagano, J. F. B., Ensink, W. A., Nehrdich, U., Spaink, H. P. and Dekker, R. J. (2020). Immunoglobulin switch-like recombination regions implicated in the formation of extrachromosomal circular 45S rDNA involved in the maternal-specific translation system of zebrafish. bioRxiv 8, 2020.01.31.928739.

Chan, K.-L., North, P. S. and Hickson, I. D. (2007). BLM is required for faithful chromosome segregation and its localization defines a class of ultrafine anaphase bridges. EMBO J 26, 3397– 3409.

Chan, Y. W., Fugger, K. and West, S. C. (2018). Unresolved recombination intermediates lead to ultra-fine anaphase bridges, chromosome breaks and aberrations. Nat. Cell Biol. 20, 92–103.

Chen, Y., Zheng, Y., Gao, Y., Lin, Z., Yang, S., Wang, T., Wang, Q., Xie, N., Hua, R., Liu, M., et al. (2018). Single-cell RNA-seq uncovers dynamic processes and critical regulators in mouse spermatogenesis. Cell Res. 28, 879–896.

Christoffels, A., Koh, E. G. L., Chia, J.-M., Brenner, S., Aparicio, S. and Venkatesh, B. (2004). Fugu genome analysis provides evidence for a whole-genome duplication early during the evolution of ray-finned fishes. Mol Biol Evol 21, 1146–1151.

Cleary, S. P., Zhang, W., Di Nicola, N., Aronson, M., Aube, J., Steinman, A., Haddad, R., Redston, M., Gallinger, S., Narod, S. A., et al. (2003). Heterozygosity for the BLM(Ash) mutation and cancer risk. Cancer Research 63, 1769–1771.

Croteau, D. L., Popuri, V., Opresko, P. L. and Bohr, V. A. (2014). Human RecQ helicases in DNA repair, recombination, and replication. Annu Rev Biochem 83, 519–552.

Cunniff, C., Bassetti, J. A. and Ellis, N. A. (2017). Bloom’s Syndrome: Clinical Spectrum, Molecular Pathogenesis, and Cancer Predisposition. MSY 8, 4–23.

de Voer, R. M., Hahn, M.-M., Mensenkamp, A. R., Hoischen, A., Gilissen, C., Henkes, A., Spruijt, L., van Zelst-Stams, W. A., Kets, C. M., Verwiel, E. T., et al. (2015). Deleterious Germline BLM Mutations and the Risk for Early-onset Colorectal Cancer. Sci Rep 5, 14060–7.

Dranow, D. B., Hu, K., Bird, A. M., Lawry, S. T., Adams, M. T., Sanchez, A., Amatruda, J. F. and Draper, B. W. (2016). Bmp15 Is an Oocyte-Produced Signal Required for Maintenance of the Adult Female Sexual Phenotype in Zebrafish. PLoS Genet 12, e1006323.

Dranow, D. B., Tucker, R. P. and Draper, B. W. (2013). Germ cells are required to maintain a stable sexual phenotype in adult zebrafish. Dev. Biol. 376, 43–50.

Ellis, N. A., Groden, J., Ye, T. Z., Straughen, J., Cell, D. L.1995 (1995). The Bloom’s syndrome gene product is homologous to RecQ helicases. Cell 83, 655–666.

Escobar, M. L., EcheverrÍa, O. M., Valenzuela, Y. M., Ortiz, R., Torres-Ramírez, N. and Vázquez-Nin, G. H. (2019). Histochemical Study of the Emergence of Apoptosis and Altered SYCP3 Protein Distribution During the First Spermatogenic Wave in Wistar Rats. Anat Rec (Hoboken) 302, 2082– 2092.

Ferencziová, V., Harami, G. M., Németh, J. B., Vellai, T. and Kovács, M. (2018). Functional fine-tuning between bacterial DNA recombination initiation and quality control systems. PLoS ONE 13, e0192483.

Gagnon, J. A., Valen, E., Thyme, S. B., Huang, P., Ahkmetova, L., Pauli, A., Montague, T. G., Zimmerman, S., Richter, C. and Schier, A. F. (2014). Efficient mutagenesis by Cas9 protein-mediated oligonucleotide insertion and large-scale assessment of single-guide RNAs. PLoS ONE 9, e98186.

German, J., Roe, A. M., Leppert, M. F. and Ellis, N. A. (1994). Bloom syndrome: an analysis of consanguineous families assigns the locus mutated to chromosome band 15q26.1. Proceedings of the National Academy of Sciences 91, 6669–6673.

Goss, K. H., Risinger, M. A., Kordich, J. J., Sanz, M. M., Straughen, J. E., Slovek, L. E., Capobianco, A. J., German, J., Boivin, G. P. and Groden, J. (2002). Enhanced tumor formation in mice heterozygous for Blm mutation. Science 297, 2051–2053.

Gruber, S. B., Ellis, N. A., Scott, K. K., Almog, R., Kolachana, P., Bonner, J. D., Kirchhoff, T., Tomsho, L. P., Nafa, K., Pierce, H., et al. (2002). BLM heterozygosity and the risk of colorectal cancer. Science 297, 2013–2013.

Hanada, K., Ukita, T., Kohno, Y., Saito, K., Kato, J. and Ikeda, H. (1997). RecQ DNA helicase is a suppressor of illegitimate recombination in Escherichia coli. Proceedings of the National Academy of Sciences 94, 3860–3865.

Hanna, C. B. and Hennebold, J. D. (2014). Ovarian germline stem cells: an unlimited source of oocytes? Fertil Steril 101, 20–30.

Harami, G. M., Seol, Y., In, J., Ferencziová, V., Martina, M., Gyimesi, M., Sarlós, K., Kovács, Z. J., Nagy, N. T., Sun, Y., et al. (2017). Shuttling along DNA and directed processing of D-loops by RecQ helicase support quality control of homologous recombination. Proc Natl Acad Sci USA 114, E466–E475.

Hatkevich, T., Kohl, K. P., McMahan, S., Hartmann, M. A., Williams, A. M. and Sekelsky, J. (2017). Bloom Syndrome Helicase Promotes Meiotic Crossover Patterning and Homolog Disjunction. Curr. Biol. 27, 96–102.

Holloway, J. K., Morelli, M. A., Borst, P. L. and Cohen, P. E. (2010). Mammalian BLM helicase is critical for integrating multiple pathways of meiotic recombination. J. Cell Biol. 188, 779–789.

Hunter, N. (2015). Meiotic Recombination: The Essence of Heredity. Cold Spring Harb Perspect Biol 7, a016618.

Karow, J. K., Chakraverty, R. K. and Hickson, I. D. (1997). The Bloom’s syndrome gene product is a 3‘-5’ DNA helicase. J. Biol. Chem. 272, 30611–30614.

Kassambara, A., Kosinski, M. and Biecek, P. (2019). survminer: Drawing Survival Curves using ‘ggplot2’. R package version 0.4.6. https://CRAN.R-project.org/package=survminer

Kossack, M. E. and Draper, B. W. (2019). Genetic regulation of sex determination and maintenance in zebrafish (Danio rerio). Curr. Top. Dev. Biol. 134, 119–149.

Krøvel, A. V. and Olsen, L. C. (2002). Expression of a vas::EGFP transgene in primordial germ cells of the zebrafish. Mech Dev 116, 141–150.

Kubota, H. and Brinster, R. L. (2018). Spermatogonial stem cells. Biol. Reprod. 99, 52–74.

Laitman, Y., Boker-Keinan, L., Berkenstadt, M., Liphsitz, I., Weissglas-Volkov, D., Ries-Levavi, L., Sarouk, I., Pras, E. and Friedman, E. (2016). The risk for developing cancer in Israeli ATM, BLM, and FANCC heterozygous mutation carriers. Cancer Genet 209, 70–74.

Leal, M. C., Cardoso, E. R., Nóbrega, R. H., Batlouni, S. R., Bogerd, J., França, L. R. and Schulz, R. W. (2009). Histological and stereological evaluation of zebrafish (Danio rerio) spermatogenesis with an emphasis on spermatogonial generations. Biol. Reprod. 81, 177–187.

Liew, W. C. and Orbán, L. (2014). Zebrafish sex: a complicated affair. Brief Funct Genomics 13, 172– 187.

Luo, Y., Hartford, S. A., Zeng, R., Southard, T. L., Shima, N. and Schimenti, J. C. (2014). Hypersensitivity of primordial germ cells to compromised replication-associated DNA repair involves ATM-p53-p21 signaling. PLoS Genet 10, e1004471.

Manthei, K. A. and Keck, J. L. (2013). The BLM dissolvasome in DNA replication and repair. Cell. Mol. Life Sci. 70, 4067–4084.

Mao, F. J., Sidorova, J. M., Lauper, J. M., Emond, M. J. and Monnat, R. J. (2010). The human WRN and BLM RecQ helicases differentially regulate cell proliferation and survival after chemotherapeutic DNA damage. Cancer Research 70, 6548–6555.

Marlow, F. (2015). Primordial Germ Cell Specification and Migration. F1000Res 4: F1000 Faculty Rev-1462.

Martin, R. H., Rademaker, A. and German, J. (1994). Chromosomal breakage in human spermatozoa, a heterozygous effect of the Bloom syndrome mutation. The American Journal of Human Genetics 55, 1242–1246.

Meeker, N. D., Hutchinson, S. A., Ho, L. and Trede, N. S. (2007). Method for isolation of PCR-ready genomic DNA from zebrafish tissues. BioTechniques 43, 610–612– 614.

O’Neil, N. J., Martin, J. S., Youds, J. L., Ward, J. D., Petalcorin, M. I. R., Rose, A. M. and Boulton, S. J. (2013). Joint molecule resolution requires the redundant activities of MUS-81 and XPF-1 during Caenorhabditis elegans meiosis. PLoS Genet 9, e1003582.

Oh, S. D., Lao, J. P., Hwang, P. Y.-H., Taylor, A. F., Smith, G. R. and Hunter, N. (2007). BLM ortholog, Sgs1, prevents aberrant crossing-over by suppressing formation of multichromatid joint molecules. Cell 130, 259–272.

Okonechnikov, K., Golosova, O., Fursov, M., and UGENE Team. (2012). Unipro UGENE: a unified bioinformatics toolkit. Bioinformatics, 28(8), 1166–1167.

Orbán, L., Sreenivasan, R. and Olsson, P.-E. (2009). Long and winding roads: testis differentiation in zebrafish. Mol. Cell. Endocrinol. 312, 35–41.

Ortega-Recalde, O., Day, R. C., Gemmell, N. J. and Hore, T. A. (2019). Zebrafish preserve global germline DNA methylation while sex-linked rDNA is amplified and demethylated during feminisation. Nat Comms 10, 3053.

Pradhan, A., Khalaf, H., Ochsner, S. A., Sreenivasan, R., Koskinen, J., Karlsson, M., Karlsson, J., McKenna, N. J., Orbán, L. and Olsson, P.-E. (2012). Activation of NF-κB protein prevents the transition from juvenile ovary to testis and promotes ovarian development in zebrafish. J. Biol. Chem. 287, 37926–37938.

Prokofyeva, D., Bogdanova, N., Dubrowinskaja, N., Bermisheva, M., Takhirova, Z., Antonenkova, N., Turmanov, N., Datsyuk, I., Gantsev, S., Christiansen, H., et al. (2013). Nonsense mutation p.Q548X in BLM, the gene mutated in Bloom’s syndrome, is associated with breast cancer in Slavic populations. Breast Cancer Res. Treat. 137, 533–539.

Ramanagoudr-Bhojappa, R., Carrington, B., Ramaswami, M., Bishop, K., Robbins, G. M., Jones, M., Harper, U., Frederickson, S. C., Kimble, D. C., Sood, R., et al. (2018). Multiplexed CRISPR/Cas9-mediated knockout of 19 Fanconi anemia pathway genes in zebrafish revealed their roles in growth, sexual development and fertility. PLoS Genet 14, e1007821.

Richardson, B. E. and Lehmann, R. (2010). Mechanisms guiding primordial germ cell migration: strategies from different organisms. Nat Rev Mol Cell Biol 11, 37–49.

Rodríguez-Marí, A. and Postlethwait, J. H. (2011). The role of Fanconi anemia/BRCA genes in zebrafish sex determination. Methods Cell Biol. 105, 461–490.

Rodríguez-Marí, A., Cañestro, C., Bremiller, R. A., Nguyen-Johnson, A., Asakawa, K., Kawakami, K. and Postlethwait, J. H. (2010). Sex reversal in zebrafish fancl mutants is caused by Tp53-mediated germ cell apoptosis. PLoS Genet 6, e1001034.

Rodríguez-Marí, A., Wilson, C., Titus, T. A., Cañestro, C., Bremiller, R. A., Yan, Y.-L., Nanda, I., Johnston, A., Kanki, J. P., Gray, E. M., et al. (2011). Roles of brca2 (fancd1) in oocyte nuclear architecture, gametogenesis, gonad tumors, and genome stability in zebrafish. PLoS Genet 7, e1001357.

Saito, D., Morinaga, C., Aoki, Y., Nakamura, S., Mitani, H., Furutani-Seiki, M., Kondoh, H. and Tanaka, M. (2007). Proliferation of germ cells during gonadal sex differentiation in medaka: Insights from germ cell-depleted mutant zenzai. Dev. Biol. 310, 280–290.

Salah, G. B., Salem, I. H., Masmoudi, A., Kallabi, F., Turki, H., Fakhfakh, F., Ayadi, H. and Kamoun, H. (2014). A novel frameshift mutation in BLM gene associated with high sister chromatid exchanges (SCE) in heterozygous family members. Mol Biol Rep 41, 7373–7380.

Sarlós, K., Biebricher, A. S., Bizard, A. H., Bakx, J. A. M., Ferreté-Bonastre, A. G., Modesti, M., Paramasivam, M., Yao, Q., Peterman, E. J. G., Wuite, G. J. L., et al. (2018). Reconstitution of anaphase DNA bridge recognition and disjunction. Nat. Struct. Mol. Biol. 25, 868–876.

Schawalder, J., Paric, E. and Neff, N. F. (2003). Telomere and ribosomal DNA repeats are chromosomal targets of the bloom syndrome DNA helicase. BMC Cell Biol. 4, 15–14.

Schayek, H., Laitman, Y., Katz, L. H., Pras, E., Ries-Levavi, L., Barak, F. and Friedman, E. (2017). Colorectal and Endometrial Cancer Risk and Age at Diagnosis in BLMAsh Mutation Carriers. Isr. Med. Assoc. J. 19, 365–367.

Schvarzstein, M., Pattabiraman, D., Libuda, D. E., Ramadugu, A., Tam, A., Martinez-Perez, E., Roelens, B., Zawadzki, K. A., Yokoo, R., Rosu, S., et al. (2014). DNA helicase HIM-6/BLM both promotes MutSγ-dependent crossovers and antagonizes MutSγ-independent interhomolog associations during caenorhabditis elegans meiosis. Genetics 198, 193–207.

Seol, Y., Harami, G. M., Kovács, M. and Neuman, K. C. (2019). Homology sensing via non-linear amplification of sequence-dependent pausing by RecQ helicase. Elife 8, 577.

Shive, H. R., West, R. R., Embree, L. J., Azuma, M., Sood, R., Liu, P. and Hickstein, D. D. (2010). brca2 in zebrafish ovarian development, spermatogenesis, and tumorigenesis. Proc Natl Acad Sci USA 107, 19350–19355.

Siegfried, K. R. and Nüsslein-Volhard, C. (2008). Germ line control of female sex determination in zebrafish. Dev. Biol. 324, 277–287.

Slanchev, K., Stebler, J., la Cueva-Méndez, de G. and Raz, E. (2005). Development without germ cells: the role of the germ line in zebrafish sex differentiation. Proceedings of the National Academy of Sciences 102, 4074–4079.

Sokolenko, A. P., Bogdanova, N., Kluzniak, W., Preobrazhenskaya, E. V., Kuligina, E. S., Iyevleva, A. G., Aleksakhina, S. N., Mitiushkina, N. V., Gorodnova, T. V., Bessonov, A. A., et al. (2014). Double heterozygotes among breast cancer patients analyzed for BRCA1, CHEK2, ATM, NBN/NBS1, and BLM germ-line mutations. Breast Cancer Res. Treat. 145, 553–562.

Sokolenko, A. P., Iyevleva, A. G., Preobrazhenskaya, E. V., Mitiushkina, N. V., Abysheva, S. N., Suspitsin, E. N., Kuligina, E. S., Gorodnova, T. V., Pfeifer, W., Togo, A. V., et al. (2012). High prevalence and breast cancer predisposing role of the BLM c.1642 C>T (Q548X) mutation in Russia. International Journal of Cancer 130, 2867–2873.

Sreenivasan, R., Jiang, J., Wang, X., Bártfai, R., Kwan, H. Y., Christoffels, A. and Orbán, L. (2014). Gonad differentiation in zebrafish is regulated by the canonical Wnt signaling pathway. Biol. Reprod. 90, 45.

Takahashi, H.1977 (1977). Juvenile hermaphroditism in the zebrafish, Brachydanio rerio. Bull. Fac. Fish. Hokkaido Univ. 28, 57–65.

Takatsu, K., Miyaoku, K., Roy, S. R., Murono, Y., Sago, T., Itagaki, H., Nakamura, M. and Tokumoto, T. (2013). Induction of female-to-male sex change in adult zebrafish by aromatase inhibitor treatment. Sci Rep 3, 3400–7.

Tangeman, L., McIlhatton, M. A., Grierson, P., Groden, J. and Acharya, S. (2016). Regulation of BLM Nucleolar Localization. Genes (Basel) 7, 69.

Taylor, J. S., Braasch, I., Frickey, T., Meyer, A. and Van de Peer, Y. (2003). Genome duplication, a trait shared by 22000 species of ray-finned fish. Genome Research 13, 382–390.

Thompson, E. R., Doyle, M. A., Ryland, G. L., Rowley, S. M., Choong, D. Y. H., Tothill, R. W., Thorne, H., kConFab, Barnes D. R., Li, J., et al. (2012). Exome sequencing identifies rare deleterious mutations in DNA repair genes FANCC and BLM as potential breast cancer susceptibility alleles. PLoS Genet 8, e1002894.

Toh, W. H., Nam, S. Y. and Sabapathy, K. (2010). An essential role for p73 in regulating mitotic cell death. Cell Death Differ 17, 787–800.

Tzung, K.-W., Goto, R., Saju, J. M., Sreenivasan, R., Saito, T., Arai, K., Yamaha, E., Hossain, M. S., Calvert, M. E. K. and Orbán, L. (2015a). Early depletion of primordial germ cells in zebrafish promotes testis formation. Stem Cell Reports 4, 61–73.

Tzung, K.-W., Goto, R., Saju, J. M., Sreenivasan, R., Saito, T., Arai, K., Yamaha, E., Hossain, M. S., Calvert, M. E. K. and Orbán, L. (2015b). Early Depletion of Primordial Germ Cells in Zebrafish Promotes Testis Formation. Stem Cell Reports 5, 156.

Uchida, D., Yamashita, M., Kitano, T. and Iguchi, T. (2002). Oocyte apoptosis during the transition from ovary-like tissue to testes during sex differentiation of juvenile zebrafish. J. Exp. Biol. 205, 711–718.

Vainio, S., Heikkilä, M., Kispert, A., Chin, N. and McMahon, A. P. (1999). Female development in mammals is regulated by Wnt-4 signalling. Nature 397, 405–409.

Walpita, D., Plug, A. W., Neff, N. F., German, J. and Ashley, T. (1999). Bloom’s syndrome protein, BLM, colocalizes with replication protein A in meiotic prophase nuclei of mammalian spermatocytes. Proceedings of the National Academy of Sciences 96, 5622–5627.

Wang, X. G., Bartfai, R., Sleptsova-Freidrich, I. and Orban, L. (2007). The timing and extent of “juvenile ovary” phase are highly variable during zebrafish testis differentiation. Journal of Fish Biology 70, 33–44.

Westerfield, M. (2000). The Zebrafish Book. 4 ed. Eugene: University of Oregon Press.

White, D. L., Mazurkiewicz, J. E. and Barrnett, R. J. (1979). A chemical mechanism for tissue staining by osmium tetroxide-ferrocyanide mixtures. J. Histochem. Cytochem. 27, 1084–1091.

White, R. J., Collins, J. E., Sealy, I. M., Wali, N., Dooley, C. M., Digby, Z., Stemple, D. L., Murphy, D. N., Billis, K., Hourlier, T., et al. (2017). A high-resolution mRNA expression time course of embryonic development in zebrafish. Elife 6, 1328.

Wickham, H. (2016). ggplot2: elegant graphics for data analysis. Springer-Verlag New York. ISBN 978-3-319-24277-4, https://ggplot2.tidyverse.org.

Wilson, C. A., High, S. K., McCluskey, B. M., Amores, A., Yan, Y.-L., Titus, T. A., Anderson, J. L., Batzel, P., Carvan, M. J., Schartl, M., et al. (2014). Wild Sex in Zebrafish: Loss of the Natural Sex Determinant in Domesticated Strains. 198, 1291–1308.

Winata, C. L., Łapiński, M., Pryszcz, L., Vaz, C., Bin Ismail, M. H., Nama, S., Hajan, H. S., Lee, S. G. P., Korzh, V., Sampath, P., et al. (2018). Cytoplasmic polyadenylation-mediated translational control of maternal mRNAs directs maternal-to-zygotic transition. Development 145, dev159566.

Wong, T.-T., Saito, T., Crodian, J. and Collodi, P. (2011). Zebrafish germline chimeras produced by transplantation of ovarian germ cells into sterile host larvae. Biol. Reprod. 84, 1190–1197.

Wu, L. and Hickson, I. D. (2003). The Bloom’s syndrome helicase suppresses crossing over during homologous recombination. Nature 426, 870–874.

Ye, D., Zhu, L., Zhang, Q., Xiong, F., Wang, H., Wang, X., He, M., Zhu, Z. and Sun, Y. (2019). Abundance of Early Embryonic Primordial Germ Cells Promotes Zebrafish Female Differentiation as Revealed by Lifetime Labeling of Germline. Mar. Biotechnol. 21, 217–228.

Yoshida, S. (2016). From cyst to tubule: innovations in vertebrate spermatogenesis. Wiley Interdiscip Rev Dev Biol 5, 119–131.

## Supplementary References

Carter, N. J., Roach, Z. A., Byrnes, M. M. and Zhu, Y. (2019). Adamts9 is necessary for ovarian development in zebrafish. Gen Comp Endocrinol 277, 130–140.

Celeghin, A., Benato, F., Pikulkaew, S., Rabbane, M. G., Colombo, L. and Dalla Valle, L. (2011). The knockdown of the maternal estrogen receptor 2a (esr2a) mRNA affects embryo transcript contents and larval development in zebrafish. Gen Comp Endocrinol 172, 120–129.

Chen, S., Zhang, H., Wang, F., Zhang, W. and Peng, G. (2016). nr0b1 (DAX1) mutation in zebrafish causes female-to-male sex reversal through abnormal gonadal proliferation and differentiation. Mol. Cell. Endocrinol. 433, 105–116.

Chen, W., Liu, L. and Ge, W. (2017). Expression analysis of growth differentiation factor 9 (Gdf9/gdf9), anti-müllerian hormone (Amh/amh) and aromatase (Cyp19a1a/cyp19a1a) during gonadal differentiation of the zebrafish, Danio rerio. Biol. Reprod. 96, 401–413.

Dai, X., Cheng, X., Huang, J., Gao, Y., Wang, D., Feng, Z., Zhai, G., Lou, Q., He, J., Wang, Z., et al. (2021). Rbm46, a novel germ cell specific factor, modulates meiotic progression and spermatogenesis. Biol. Reprod.

Feng, K., Cui, X., Song, Y., Tao, B., Chen, J., Wang, J., Liu, S., Sun, Y., Zhu, Z., Trudeau, V. L., et al. (2020). Gnrh3 Regulates PGC Proliferation and Sex Differentiation in Developing Zebrafish. Endocrinology 161.

Goudarzi, M., Banisch, T. U., Mobin, M. B., Maghelli, N., Tarbashevich, K., Strate, I., van den Berg, J., Blaser, H., Bandemer, S., Paluch, E., et al. (2012). Identification and regulation of a molecular module for bleb-based cell motility. Dev Cell 23, 210–218.

Hartung, O., Forbes, M. M. and Marlow, F. L. (2014). Zebrafish vasa is required for germ-cell differentiation and maintenance. Mol Reprod Dev 81, 946–961.

Houwing, S., Berezikov, E. and Ketting, R. F. (2008). Zili is required for germ cell differentiation and meiosis in zebrafish. EMBO J 27, 2702–2711.

Houwing, S., Kamminga, L. M., Berezikov, E., Cronembold, D., Girard, A., van den Elst, H., Filippov, D. V., Blaser, H., Raz, E., Moens, C. B., et al. (2007). A role for Piwi and piRNAs in germ cell maintenance and transposon silencing in Zebrafish. Cell 129, 69–82.

Kaufman, O. H., Lee, K., Martin, M., Rothhämel, S. and Marlow, F. L. (2018). rbpms2 functions in Balbiani body architecture and ovary fate. PLoS Genet 14, e1007489.

Kossack, M. E., High, S. K., Hopton, R. E., Yan, Y.-L., Postlethwait, J. H. and Draper, B. W. (2019). Female Sex Development and Reproductive Duct Formation Depend on Wnt4a in Zebrafish. Genetics 211, 219–233.

Lau, E. S.-W., Zhang, Z., Qin, M. and Ge, W. (2016). Knockout of Zebrafish Ovarian Aromatase Gene (cyp19a1a) by TALEN and CRISPR/Cas9 Leads to All-male Offspring Due to Failed Ovarian Differentiation. Sci Rep 6, 37357.

Leerberg, D. M., Sano, K. and Draper, B. W. (2017). Fibroblast growth factor signaling is required for early somatic gonad development in zebrafish. PLoS Genet 13, e1006993.

Lu, H., Cui, Y., Jiang, L. and Ge, W. (2017). Functional Analysis of Nuclear Estrogen Receptors in Zebrafish Reproduction by Genome Editing Approach. Endocrinology 158, 2292–2308.

Miao, L., Yuan, Y., Cheng, F., Fang, J., Zhou, F., Ma, W., Jiang, Y., Huang, X., Wang, Y., Shan, L., et al. (2017). Translation repression by maternal RNA binding protein Zar1 is essential for early oogenesis in zebrafish. Development 144, 128–138.

Qin, M., Zhang, Z., Song, W., Wong, Q. W.-L., Chen, W., Shirgaonkar, N. and Ge, W. (2018). Roles of Figla/figla in Juvenile Ovary Development and Follicle Formation During Zebrafish Gonadogenesis. Endocrinology 159, 3699–3722.

Romano, S., Kaufman, O. H. and Marlow, F. L. (2020). Loss of dmrt1 restores zebrafish female fates in the absence of cyp19a1a but not rbpms2a/b. Development 147.

Sone, R., Taimatsu, K., Ohga, R., Nishimura, T., Tanaka, M. and Kawahara, A. (2020). Critical roles of the ddx5 gene in zebrafish sex differentiation and oocyte maturation. Sci Rep 10, 14157.

Vong, Y. H., Sivashanmugam, L., Zaucker, A., Jones, A. and Sampath, K. (2020). The RNA-binding protein Igf2bp3 is critical for embryonic and germline development in zebrafish. bioRxiv 2020.06.23.167163.

Weidinger, G., Stebler, J., Slanchev, K., Dumstrei, K., Wise, C., Lovell-Badge, R., Thisse, C., Thisse, B. and Raz, E. (2003). dead end, a novel vertebrate germ plasm component, is required for zebrafish primordial germ cell migration and survival. Curr. Biol. 13, 1429–1434.

Xia, H., Zhong, C., Wu, X., Chen, J., Tao, B., Xia, X., Shi, M., Zhu, Z., Trudeau, V. L. and Hu, W. (2018). Mettl3 Mutation Disrupts Gamete Maturation and Reduces Fertility in Zebrafish. Genetics 208, 729–743.

Xie, Y., Huang, D., Chu, L., Liu, Y., Sun, X., Li, J. and Cheng, C. H. K. (2020). Igf3 is essential for ovary differentiation in zebrafish. Biol. Reprod.

Yang, Y.-J., Wang, Y., Li, Z., Zhou, L. and Gui, J.-F. (2017). Sequential, Divergent, and Cooperative Requirements of Foxl2a and Foxl2b in Ovary Development and Maintenance of Zebrafish. Genetics 205, 1551–1572.

Yin, Y., Tang, H., Liu, Y., Chen, Y., Li, G., Liu, X. and Lin, H. (2017). Targeted Disruption of Aromatase Reveals Dual Functions of cyp19a1a During Sex Differentiation in Zebrafish. Endocrinology 158, 3030–3041.

Zhang, Z., Lau, S.-W., Zhang, L. and Ge, W. (2015a). Disruption of Zebrafish Follicle-Stimulating Hormone Receptor (fshr) But Not Luteinizing Hormone Receptor (lhcgr) Gene by TALEN Leads to Failed Follicle Activation in Females Followed by Sexual Reversal to Males. Endocrinology 156, 3747–3762.

Zhang, Z., Zhu, B. and Ge, W. (2015b). Genetic analysis of zebrafish gonadotropin (FSH and LH) functions by TALEN-mediated gene disruption. Mol. Endocrinol. 29, 76–98.

Zhu, J., Zhang, D., Liu, X., Yu, G., Cai, X., Xu, C., Rong, F., Ouyang, G., Wang, J. and Xiao, W. (2019). Zebrafish prmt5 arginine methyltransferase is essential for germ cell development. Development 146.

